# RNA structural dynamics modulate EGFR-TKIs resistance through controlling *YRDC* translation in NSCLC cells

**DOI:** 10.1101/2022.10.17.512459

**Authors:** Boyang Shi, Ke An, Yueqin Wang, Yuhan Fei, Caixia Guo, Qiangfeng Cliff Zhang, Yun-gui Yang, Xin Tian, Quancheng Kan

**Author notes:** These authors contributed equally (Boyang Shi, Ke An).

## Abstract

Epidermal growth factor receptor-tyrosine kinase inhibitors (EGFR-TKIs) positively affect the initial control of non-small cell lung cancer (NSCLC). The rapidly acquired TKIs resistance accounts for a major hurdle in successful treatment. However, the mechanisms controlling EGFR-TKIs resistance remain largely unknown. RNA structures have widespread and crucial roles in various biological processes; but, their role in regulating cancer drug resistance remains unclear. Here, the PARIS method is used to establish the higher-order RNA structure maps of EGFR-TKI resistant- and sensitive-cells of NSCLC. According to our results, RNA structural regions are enriched in UTRs and correlate with translation efficiency. Moreover, *YRDC* facilitates resistance to EGFR-TKIs in NSCLC cells, and RNA structure formation in *YRDC* 3’UTR suppress ELAVL1 binding leading to EGFR-TKIs sensitivity by impairing *YRDC* translation. A potential cancer therapy strategy is provided by using antisense oligonucleotide (ASO) to perturb the interaction between RNA and protein. Our study reveals an unprecedented mechanism in which the RNA structure switch modulates EGFR-TKIs resistance by controlling *YRDC* mRNA translation in an ELAVL1-dependent manner.

## INTRODUCTION

Lung cancer (LC) is a crucial factor in cancer-related mortality worldwide, regardless of gender. Non-small cell lung cancer (NSCLC) accounts for a primary LC histological subtype, constituting 85% of all lung cancer cases(Siegel et al., 2021). Although conventional cytotoxic chemotherapy is still a critical method to treat advanced NSCLC, the latest advancements in individualized medicine have enhanced the responsiveness of oncogenic mutation-harboring NSCLC patients to targeted treatment without inducing severe side effects. Particularly, EGFR-specific tyrosine kinase inhibitors (TKIs) are widely studied(Hirsch et al., 2017), and *EGFR* genetic mutations are usually present among NSCLC cases (incidence of >50%)(Shi et al., 2014). NSCLC patients harboring these sensitizing EGFR mutations were exceptionally sensitive to the reversible first-generation EGFR TKIs (gefitinib and erlotinib) and the irreversible second-generation EGFR TKIs (afatinib and dacomitinib)(Piotrowska and Sequist, 2016; Wu and Fu, 2018). Although disease control and clinical responsiveness rates attained with EGFR-TKIs were excellent, resistance soon developed in such cases (mean, 1-year)(Yu et al., 2013). Among the various resistance mechanisms of EGFR-TKIs, EGFR gatekeeper T790M point mutation near the catalytic site has the highest prevalence. EGFR T790M mutation elevates ATP affinity in receptor tyrosine kinase and shows steric hindrance of EGFR-TKI binding, resulting in TKI treatment failure(Recondo et al., 2018). Although third-generation EGFR-TKIs (AZD9291, CO-1686, and HM61713) could overcome EGFR mutation threonine 790 (T790M) resistance and are becoming the new first-line standard in EGFR mutant NSCLC, acquired resistance is virtually inevitable(Sullivan and Planchard, 2016), indicating these genetic events are insufficient to explain TKI resistance. Consequently, it is crucial and urgent to identify novel therapeutics or treatments to treat LC with EGFR-TKI resistance.

RNA molecules can fold into complicated structures necessary for the diverse roles and regulation, such as transcription, splicing, polyadenylation, degradation, translation, and localization in cells(Ganser et al., 2019; Lewis et al., 2017; Mustoe et al., 2018; Wang et al., 2021b). Recently, couple of chemical probing with high-throughput sequencing methods have been used to study whole-transcriptome secondary structures. Whole-genome RNA secondary structural probing with dimethylsulfate-sequencing (DMS-seq)(Ding et al., 2014; Rouskin et al., 2014), and SHAPE-sequencing(Smola et al., 2015; Spitale et al., 2015) is conducted in live cells, which revealed the active mRNA structural unfolding, indicating the contribution of RNA structures to its overall processing. The above approaches indicated significant progress and offered specific data on the single-or double-stranded RNA regions in cells but did not detect the direct pairing information between RNA sequences. By crosslinking the base-paired RNAs to psoralen under UV irradiation(Aw et al., 2016; Graveley, 2016; Sharma et al., 2016), PARIS (Psoralen Analysis of RNA Interactions and Structures) could identify the detailed intermolecular or intramolecular base-pairing pattern *in vivo*(Lu et al., 2016).

Over the past few years, many studies revealed that the widespread regulation of RNA processing, including post-transcriptional regulation in cancer impacts multiple facets of tumorigenesis and drug resistance(Delaunay and Frye, 2019; Goodall and Wickramasinghe, 2021). Meanwhile, RNA structures are crucial in physiological processes, including embryogenesis(Beaudoin et al., 2018; Shi et al., 2020), cardiac specification(Xue et al., 2016), neurogenesis(Bernat and Disney, 2015; Wang et al., 2021a), viral infection(Cao et al., 2021; Huston et al., 2021; Mizrahi et al., 2018; Sun et al., 2021; Ziv et al., 2020). Consequently, regulation based on RNA structures is expected to affect tumorigenesis and drug resistance critically. Our study illustrated the *in vivo* RNA structural landscapes in cells with EGFR-TKIs resistance and sensitivity using PARIS, and found that the RNA structure switch modulates EGFR-TKIs resistance by regulating *YRDC* mRNA translation in an ELAVL1-dependent manner. Collectively, we demonstrated a previously unappreciated yet fundamental role of RNA structure-based regulation of EGFR-TKIs resistance.

## RESULTS

### Higher-Order RNA Structure maps of cells with EGFR-TKI resistance and sensitivity

The AZD9291-resistant (AZD9291-R) (Figure 1A) and gefitinib-resistant (Gefitinib-R) cells (Figure 1B) were established using EGFR-mutant NSCLC cell line (PC9) to explore the mechanisms of RNA structures in lung cancer acquired EGFR-TKIs resistance. *In-situ* PARIS assay(Lu et al., 2016), a high-throughput sequencing methods technology that allows the potent measurement of RNA duplexes that show near base-pair resolution in PC9, PC9(Gefitinib-R), or PC9(AZD9291-R) cells, was performed (Figure 1C). We determined the RNA structures as previously described(Lu et al., 2016) and obtained transcriptome-wide RNA structural maps with a high correlation between biological replicates (Figures S1A-D, Table S1). Overall, about 150,000 RNA duplexes from more than 15,000 transcripts were obtained for each of the cell lines (Figure 1D), including intramolecular (Figure S1E) and intermolecular RNA duplexes (Figure S1F). Our study reveals that RNA structures involve most functional classes of RNAs, including mRNA, miRNA, lncRNA, snoRNA, and snRNA (Figure 1E, Figures S1G, H). Overall, our study constructed RNA structure maps of cells with EGFR-TKI resistance and sensitivity at the transcriptome level.

**Figure 1.**
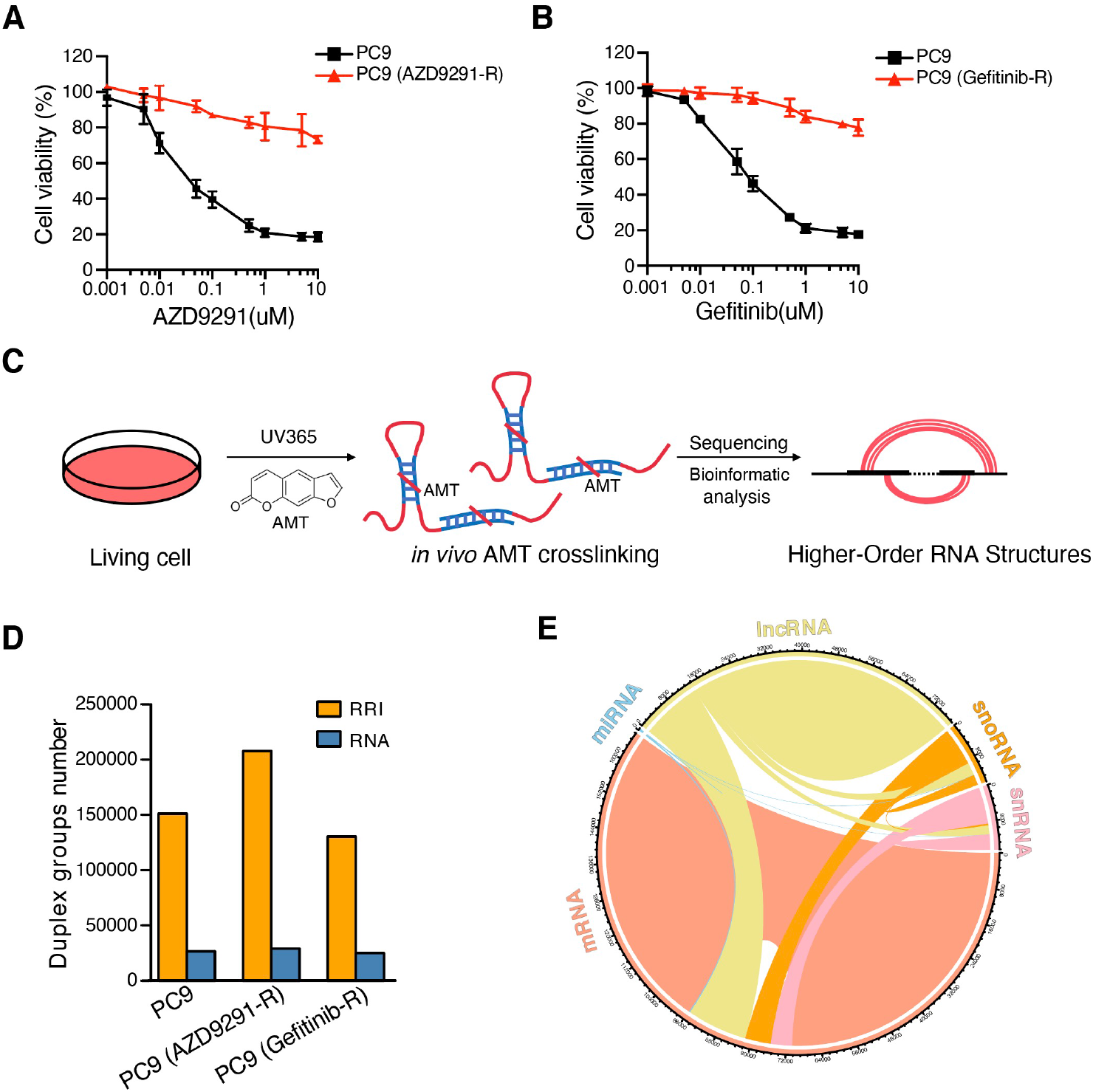
Comprehensive Analysis of RNA Structures in EGFR-TKIs Resistant- and Sensitive-cells. **(A,B)** CCK-8 assays in PC9, PC9 (AZD9291-R) or PC9 (Gefitinib-R) cells treated with AZD9291 (**A**) or Gefitinib (**B**). **(C)** Schematic view of *in vivo* RNA structures maps in TKIs resistant- and sensitive-cells using PARIS. **(D)** The number of RNA-RNA duplexes and transcripts in EGFR-TKIs resistant- and sensitive-cells. **(E)** Circos plot of the landscape of RNA-RNA duplexes detected by PARIS in PC9 cells.

Having established the RNA structure maps of cells with EGFR-TKI resistance and sensitivity, bioinformatic analyses were performed to investigate the RNA structure’s global characteristics. Unlike the previous study which can only identify local structures, mainly within the window of <200 nt, our study identified numerous RNA duplexes (more than 50%) spanning>200 nt, with more than 30% of them spanning over 1,000 nt in cells with EGFR-TKI resistance and sensitivity (Figure 2A). To study the RNA structural distribution of mRNA, the two-dimensional heatmaps of enriched RNA structural sites down the mRNA length that aligned transcripts in line with the sites of translation initiation and termination codons were plotted (Figure 2B, Figures S2A, B). Intriguingly, we found that RNA structures were also enriched in UTR, translation initiation sites, and stop codons. Except for the local RNA structures, there were many long-range RNA structures across different regions, especially in the UTR. UTR is crucial for RNA processing regulation. Thus, RNA structures are the possible regulator for RNA processing modulation in cells with EGFR-TKI resistance and sensitivity.

**Figure 2.**
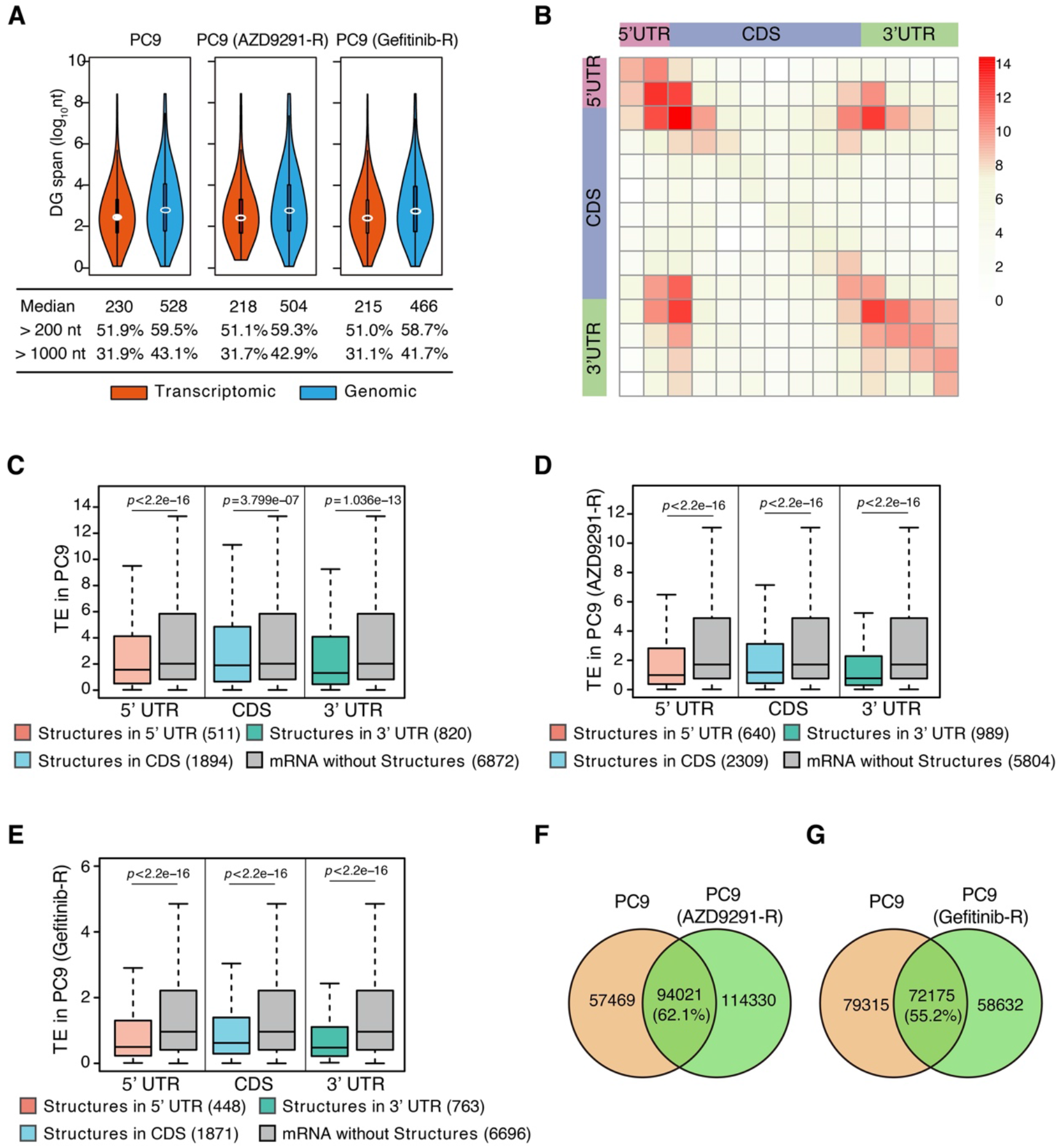
Genome-wide Analysis of Features in RNA Structures in EGFR-TKIs Resistant- and Sensitive-cells. **(A)** Size distribution of RNA duplexes in transcriptome and genome. Genomic span is the distance between the ends of gapped reads in the genome, while the transcriptomic span excludes introns. **(B)** Two-dimensional heatmap showing enrichment of mRNA duplexes based on the location of chimera ends in PC9 cells. **(C-E)** Boxplot chart showing decreased translation efficiency (TE) for EGFR-TKIs sensitive- and resistant-cells mRNAs displaying RNA structures within 5’UTR, CDS or 3’UTR compared to mRNAs without RNA structures. *P* values were calculated by the Wilcox. test. **(F)** Overlay of RNA duplex groups between PC9 cells and PC9(AZD9291-R) cells. **(G)** Overlay of RNA duplex groups between PC9 cells and PC9(Gefitinib-R) cells.

To explore whether RNA structures are involved in RNA translation, the ribosome profiling assay and RNA-seq in PC9, PC9 (AZD9291-R), or PC9 (Gefitinib-R) cells were performed (Figures S3A-C). Intriguingly, for structural mRNAs, their translation efficiency (TE) experienced significant impairment compared to non-structural counterparts in these three cell types (Figures S3D). Furthermore, we examined the translation efficiency of mRNAs with RNA structures located in different regions (5’ UTR, CDS, and 3’ UTR), and found that RNA structures in all regions impaired mRNA translation efficiency, compared to the non-structural mRNA in all three cell lines (Figures 2C-E). Collectively, these results suggest that conserved regulation of RNA structure on translation in cells with EGFR-TKI resistance and sensitivity. To further evaluate RNA structure changes in the transition of EGFR-TKIs resistance, the RNA duplexes were compared between cells with EGFR-TKI resistance and sensitivity. Almost 60% of RNA duplexes were conserved between cells with EGFR-TKI resistance and sensitivity (Figures 2F, G), indicating the dynamic RNA structures in the transition of EGFR-TKIs resistance. In sum, the RNA structural changes between cells with EGFR-TKI resistance and sensitivity are not found at the whole transcriptome level. Considering that drug resistance arises from the evolutionary pressure exerted on cancer cells(Jamal-Hanjani et al., 2017; Vasan et al., 2019), we speculate that the changes in RNA structural regulations mainly occur at the transcripts level.

### *YRDC* facilitates LC cell resistance to EGFR-TKIs

To determine whether the translation regulation is a crucial regulator for EGFR-TKIs resistance, the translation efficiency of cells with EGFR-TKI resistance relative to those with sensitivity was compared. As a result, EGFR-TKIs resistance cause translation changes in both PC9 (Gefitinib-R) and PC9 (AZD9291-R) cells compared to PC9 cells (Figure 3A). Intriguingly, an 80.1% (1766/(1766+437)) overlap in PC9 (Gefitinib-R) cells was observed between the upregulated genes in these two EGFR-TKI resistant cells, and 93.7% (6524/(6524+437)) overlap in PC9 (Gefitinib-R) cells between the downregulated genes (Figure 3A). Hence, the regulation of translation between EGFR-TKI resistant and sensitive cells exhibits the similar tendency on translation efficiency of these changed genes. Based on the above findings, translation control is speculated to regulate EGFR-TKIs resistance. Gene Ontology (GO) analysis showed that common upregulated genes in EGFR-TKI resistant cells are enriched in processes of intracellular signal transduction, MAPK cascade, Ras protein signal transduction, regulation of Wnt signaling pathway, and small GTPase mediated signal transduction (Figure 3B, Table S2). Common downregulated genes in EGFR-TKI resistant cells are enriched in processes of DNA repair, cell cycle, cellular response to DNA damage stimulus, and response to drug (Figure 3C, Table S2). These results suggested that the translation changes may play a crucial role in EGFR-TKI resistance.

**Figure 3.**
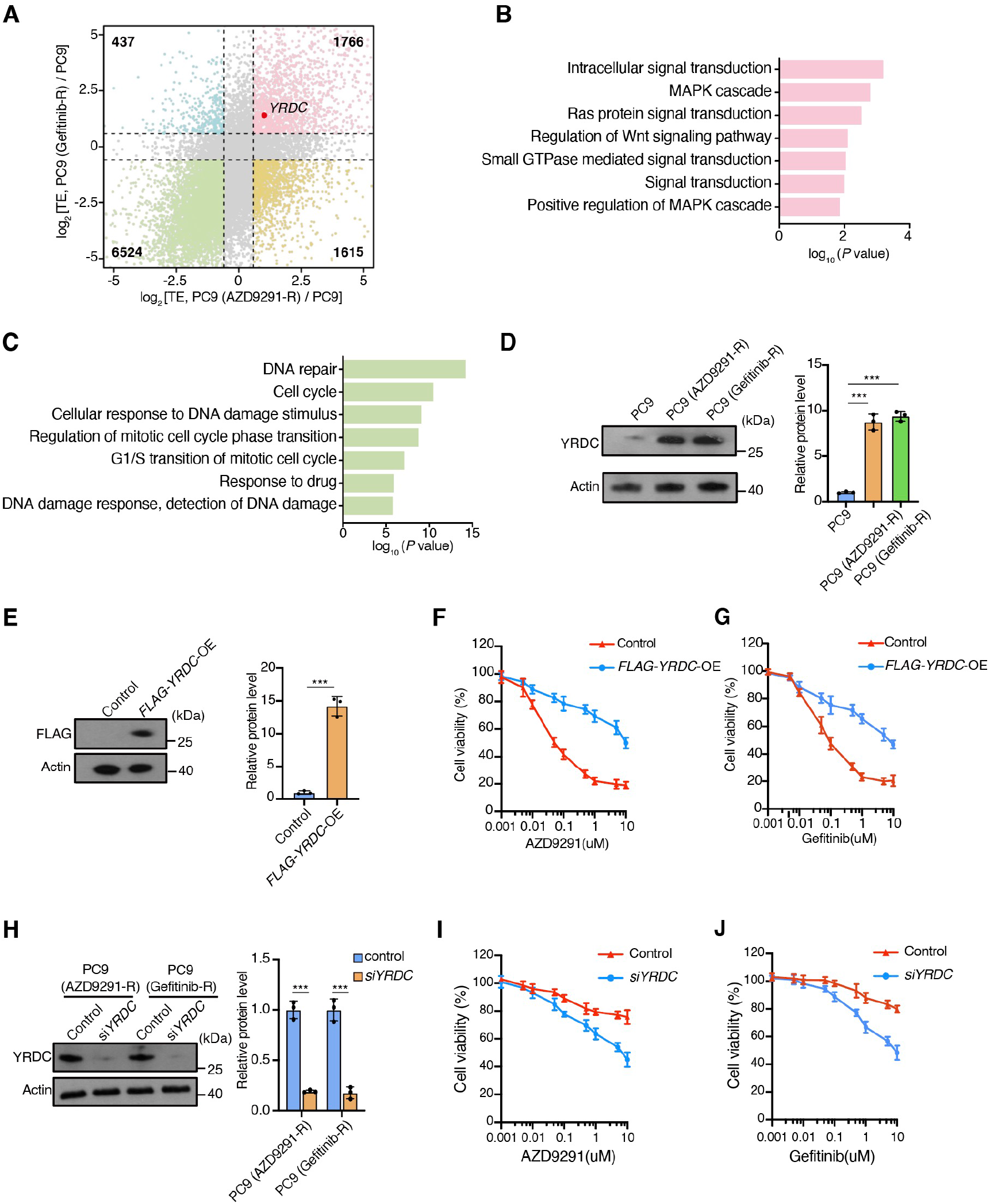
*YRDC* is Required for EGFR-TKIs Resistance. **(A)** Volcano plots showing the log2 fold changes of TE in PC9(AZD9291-R) cells or PC9(Gefitinib-R) cells versus PC9 cells. **(B)** Gene set enrichment analysis of common upregulated genes compared between EGFR-TKI resistant- and sensitive-cells. **(C)** Gene set enrichment analysis of common downregulated genes compared between EGFR-TKI resistant- and sensitive-cells. **(D)** Western blot (left) and statistical (right) analyses show protein level of YRDC in PC9 cells, PC9(AZD9291-R) cells and PC9(Gefitinib-R) cells. n = 3 biological replicates, error bars, mean ± SD, *P* values were calculated by the two-tailed Student’ s *t*-test, ***, *P* < 0.001. **(E)** Western blot (left) and statistical (right) analyses show protein level of control and *FLAG-YRDC* overexpression in PC9 cells. n = 3 biological replicates, error bars, mean ± SD, *P* values were calculated by the two-tailed Student’ s *t*-test, ***, *P* < 0.001. **(F,G)** CCK-8 assays for PC9 cells transfected with *FLAG-YRDC* expression or control vectors for 24 h followed by AZD9291 (**F**) or Gefitinib (**G**) treatment for another 48 h. **(H)** Western blot (left) and statistical (right) analyses show protein level of control and *YRDC* knockdown in PC9 cells. n = 3 biological replicates, error bars, mean ± SD, *P* values were calculated by the two-tailed Student’ s *t*-test, ***, *P* < 0.001. **(I,J)** CCK-8 assays for PC9(AZD9291-R) (**I**) cells and PC9(Gefitinib-R) (**J**) with or without *YRDC* knockdown for 24 h followed by AZD9291 (**I**) or Gefitinib (**J**) treatment for another 48 h.

Previous studies have reported that *YRDC* regulates RNA translation via involvement in tRNA’s N6-threonyl carbamoyl adenosine (t^6^A) synthesis(El Yacoubi et al., 2009; Lescrinier et al., 2006; Murphy et al., 2004), and it has been reported that can regulate HCC cell resistance to lenvatinib by regulating KRAS translation(Guo et al., 2021). In our study, we found that *YRDC* has a higher translation efficiency in EGFR-TKI resistant cells than sensitive cells (Figure 3A). YRDC protein expression was verified through Western blot, which showed high expression in EGFR-TKI resistant cells (Figure 3D). We also investigated the aberrant expression of *YRDC* mRNA in the TCGA lung cancer cohort and observed markedly increased *YRDC* expression in NSCLC specimens compared to normal tissue (Figure S4A), indicating its role in tumorigenesis. Thus, we speculate that *YRDC* might be involved in regulating NSCLC cell resistance to EGFR-TKIs. The CDS region of *YRDC* overexpression by plasmid transfection in EGFR-TKI sensitive cells (PC9 cells) (Figure 3E) was performed to test this hypothesis. Further, a CCK8 assay was conducted to detect the cell viability under AZD9291 and Gefitinib’s respective treatments. *YRDC* overexpression induced a higher EGFR-TKIs resistance than the control group in PC9 cells (Figures 3F, G). *YRDC* knockdown by siRNA transfection into cells with EGFR-TKI resistance (PC9 (Gefitinib-R) and PC9 (AZD9291-R)) was also performed (Figure 3H). The resistance to EGFR-TKIs was impaired under *YRDC* knockdown (Figures 3I, J), indicating that *YRDC* facilitates NSCLC cells resistance to EGFR-TKIs.

Intriguingly, by analyzing PARIS data, we found that 3’ UTR of *YRDC* mRNA forms a double-strand structure only in EGFR-TKI sensitive cells (PC9) (Figure S4B), which unwound in both EGFR-TKI resistant cells (PC9 (Gefitinib-R) and PC9 (AZD9291-R)). Our results have suggested that the RNA structures impair the translation efficiency in cells with EGFR-TKI resistance and sensitivity; thus, we speculate that the structural changes may influence the translation of *YRDC* mRNA.

### RNA structure in *YRDC* 3’ UTR is necessary for EGFR-TKIs resistance

For investigating RNA structure’s effect on *YRDC* 3’ UTR for translation control and EGFR-TKIs resistance, four mutants containing endogenous CDS and 3’ UTR of *YRDC* with point mutations (Mut-1: U1577C+G1582C, Mut-2: G1585C+G1590U, Mut-3: U1631G+A1636C, Mut-4: C1625A+U1628G) that disrupt base pairing (Figure 4A, Figure S5) were constructed and transfected PC9 cells with wild-type or mutant *YRDC* plasmids. Only the Mut-1 group showed increased protein expression level (Figure 4B) from the Western-blot assay; however, mRNA expression was not changed significantly between wild-type or mutants (Figure 4C). As a result, RNA structure in *YRDC* 3’ UTR impairs the *YRDC* protein translation, not the mRNA abundance. Meanwhile, only Mut-1 abolishes this RNA structure’s translation inhibition, indicating the stem formed by 1575–1584 nt and 1629–1638 nt of *YRDC* mRNA was essential for translation control. Furthermore, Mut-1 and Mut-3 can disrupt the base pairing of 1575– 1584 nt and 1629–1638 nt of *YRDC* mRNA. However, only Mut-1 (mutant position at 1575–1584 nt of *YRDC* mRNA) abolished the translation inhibition, but not *Mut-3* (mutant position at 1629–1637 nt of *YRDC* mRNA), indicating that the region of 1629– 1637 nt is a functional region that modulates translation of 1629–1637 nt of *YRDC* mRNA. The region of 1575–1584 nt may play the role of RNA structural regulator as a flanking sequence. The results of the CCK8 assay also showed that only Mut-1 overexpression induces a higher resistance to EGFR-TKIs (Figures 4D, E), supporting that the RNA structure in *YRDC* mRNA 3’ UTR possibly has an essential effect on translation control and EGFR-TKIs resistance. Antisense oligonucleotide (ASO) was also used to modulate the RNA structure in *YRDC* 3’ UTR in EGFR-TKIs resistant cells, to test whether this RNA structure can influence EGFR-TKIs resistance. ASO was prepared with the phosphorothioate backbone and 2’-O-methoxyethyl (2’-MOE) modifications to reduce cell toxicity and enhance nuclease resistance. The results showed that ASO-YRDC (antisense of 1624–1643 nt in *YRDC* mRNA) transfection groups had a lower YRDC protein expression than ASO-NC (non-binding ASO) groups in EGFT-TKIs resistant cells (Figures 4F, I). Besides, the mRNA expression level was not influenced by ASOs transfection (Figures 4G, J). CCK8 assay showed that ASO-YRDC transfection decreased the resistance to EGFR-TKIs (Figures 4H, K), indicating that the RNA structure modulation in EGFR-TKIs resistance cells by ASO transfection can restore the sensitivity to EGFR-TKIs.

**Figure 4.**
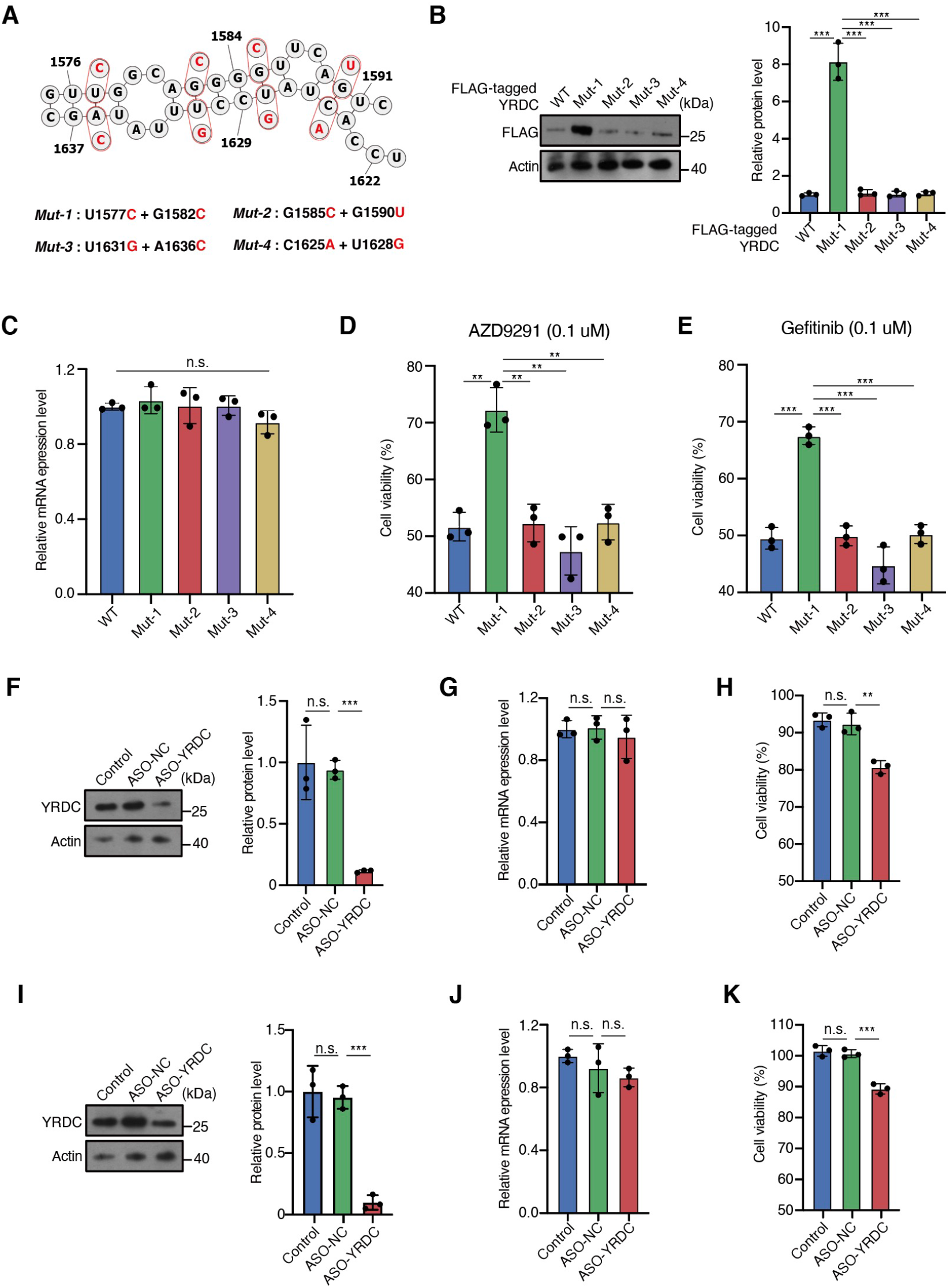
RNA Structure in 3’ UTR Region of *YRDC* Contributes to EGFR-TKIs Resistance via Controlling *YRDC* mRNA Translation. **(A)** Predicted secondary structure model of the RNA structure in 3’ UTR region of *YRDC* mRNA, annotated with genomic coordinates. Red circles represent designs of mutations in the transfection study. **(B)** Western blot (left) and statistical (right) analyses show protein level of FLAG-YRDC or different FLAG-YRDC mutations overexpression in PC9 cells. n = 3 biological replicates, error bars, mean ± SD, *P* values were calculated by the two-tailed Student’ s *t*-test, ***, *P* < 0.001. **(C)** Relative mRNA level of the *FLAG-YRDC* or different *FLAG-YRDC* mutations reporter gene in PC9 cells. n = 3, error bars, mean ± SD, *P* values were determined by the two-tailed Student’ s *t*-test, n.s., *P*, not significant. **(D,E)** CCK-8 assays for PC9 cells transfected with *FLAG-YRDC* or different *FLAG-YRDC* mutations vectors for 24 h followed by AZD9291 (0.1uM) (**D**) or Gefitinib (0.1uM) (**E**) treatment for another 48 h. n = 3, error bars, mean ± SD, *P* values were determined by the two-tailed Student’ s *t*-test, **, *P* < 0.01, ***, *P* < 0.001. **(F)** Western blot (left) and statistical (right) analyses show protein level of YRDC in PC9(AZD9291-R) cells in control, ASO-NC or ASO-YRDC transfection groups. n = 3 biological replicates, error bars, mean ± SD, *P* values were calculated by the two-tailed Student’ s *t*-test, ***, *P* < 0.001, n.s., *P*, not significant. **(G)** Relative mRNA level of the *YRDC* in PC9(AZD9291-R) cells in control, ASO-NC or ASO-YRDC transfection groups. n = 3, error bars, mean ± SD, *P* values were determined by the two-tailed Student’ s *t*-test, n.s., *P*, not significant. **(H)** CCK-8 assays for PC9(AZD9291-R) cells transfected with ASO-NC or ASO-YRDC for 24 h followed by AZD9291 (0.1uM) treatment for another 48 h. n = 3 biological replicates, error bars, mean ± SD, *P* values were calculated by the two-tailed Student’ s *t*-test, **, *P* < 0.01, n.s., *P*, not significant. **(I)** Western blot (left) and statistical (right) analyses show protein level of YRDC in PC9(Gefitinib-R) cells in control, ASO-NC or ASO-YRDC transfection groups. n = 3 biological replicates, error bars, mean ± SD, *P* values were calculated by the two-tailed Student’ s *t*-test, ***, *P* < 0.001, n.s., *P*, not significant. **(J)** Relative mRNA level of the *YRDC* in PC9(Gefitinib-R) cells in control, ASO-NC or ASO-YRDC transfection groups. n = 3, error bars, mean ± SD, *P* values were determined by the two-tailed Student’ s *t*-test,, n.s., *P*, not significant. **(K)** CCK-8 assays for PC9(Gefitinib-R) cells transfected with ASO-NC or ASO-YRDC for 24 h followed by Gefitinib (0.1uM) treatment for another 48 h, n = 3 biological replicates, error bars, mean ± SD, *P* values were calculated by the two-tailed Student’ s *t*-test, ***, *P* < 0.001, n.s., *P*, not significant.

### ELAVL1 shows a higher affinity for single-stranded RNA within 3’ UTR of *YRDC in-vitro* and *in-vivo*

According to previous studies, RNA structures can regulate RBPs (RNA binding proteins) binding to RNA by structural switching(Shi et al., 2020; Spitale et al., 2015). Therefore, RNA structure in *YRDC* mRNA 3’ UTR might regulate translation by modulating RBPs binding. By analyzing the sequencing of 1629–1637 nt of *YRDC* mRNA, we found that this region contains the sequence of UUUAUA, an AU-rich element (ARE). ARE is an important *cis*-element for RNA processing, and previous study have demonstrated that ARE could regulate ELAVL1 binding by RNA structure switch(Shi et al., 2020). ELAVL1 is well-known for regulating RNA stability and translation(Hinman and Lou, 2008). Thus, we speculated that RNA structure in *YRDC* 3’ UTR regulates *YRDC* mRNA translation by controlling ELAVL1 protein binding. Flag-ELAVL1 RIP-qPCR was performed to test the ELAVL1 binding on *YRDC*. The results showed that ELAVL1 could bind *YRDC* in PC9 (AZD9291-R) and PC9 (Gefitinib-R), but not PC9 cells (Figure 5A, Figure S6A). To further examine the binding ability of ELAVL1, biotin-labeled RNA pulldown assays were conducted using an RNA probe of 1621–1638 nt of *YRDC* mRNA. The results of both in vivo and in vitro RNA pulldown assays demonstrated that the ELAVL1 protein can be more efficiently pulled down by the RNA probe, comparing the negative control (Figures 5B, C). Similarly, electrophoretic mobility shift assays (EMSA) (Figure S6B) also illustrated that ELAVL1 could bind the region of 1621–1638 nt of *YRDC* mRNA. Considering the RNA structure switch between cells with EGFR-TKI resistance and those with sensitivity, these results hinted that RNA structure in *YRDC* 3’ UTR regulates ELAVL1 binding in a structure-dependent manner. For testing the role of RNA structure in ELAVL1’s binding on 3’ UTR of *YRDC* mRNA, RNA probes were synthesized to reconstruct the wild type RNA structure formed by 1575–1592 nt and 1621-1638 nt of *YRDC* mRNA (Wild type), Rescue and Mutant with restored and disrupted base pairing within ARE region, respectively (Figure 5D). These assays were repeated by adopting diverse biotin-labeled probe structures. According to *in-vitro* and *in-vivo* pulldown (Figures 5E, F) and EMSA (Figure 5G) results, ELAVL1 preferentially bound to 3’ UTR of *YRDC* mRNA with single-strand than the double-strand. Overall, the results indicate that the 3’ UTR of *YRDC* mRNA could efficiently affect ELAVL1 protein binding by switching RNA structure between EGFR-TKI resistant and sensitive cells.

**Figure 5.**
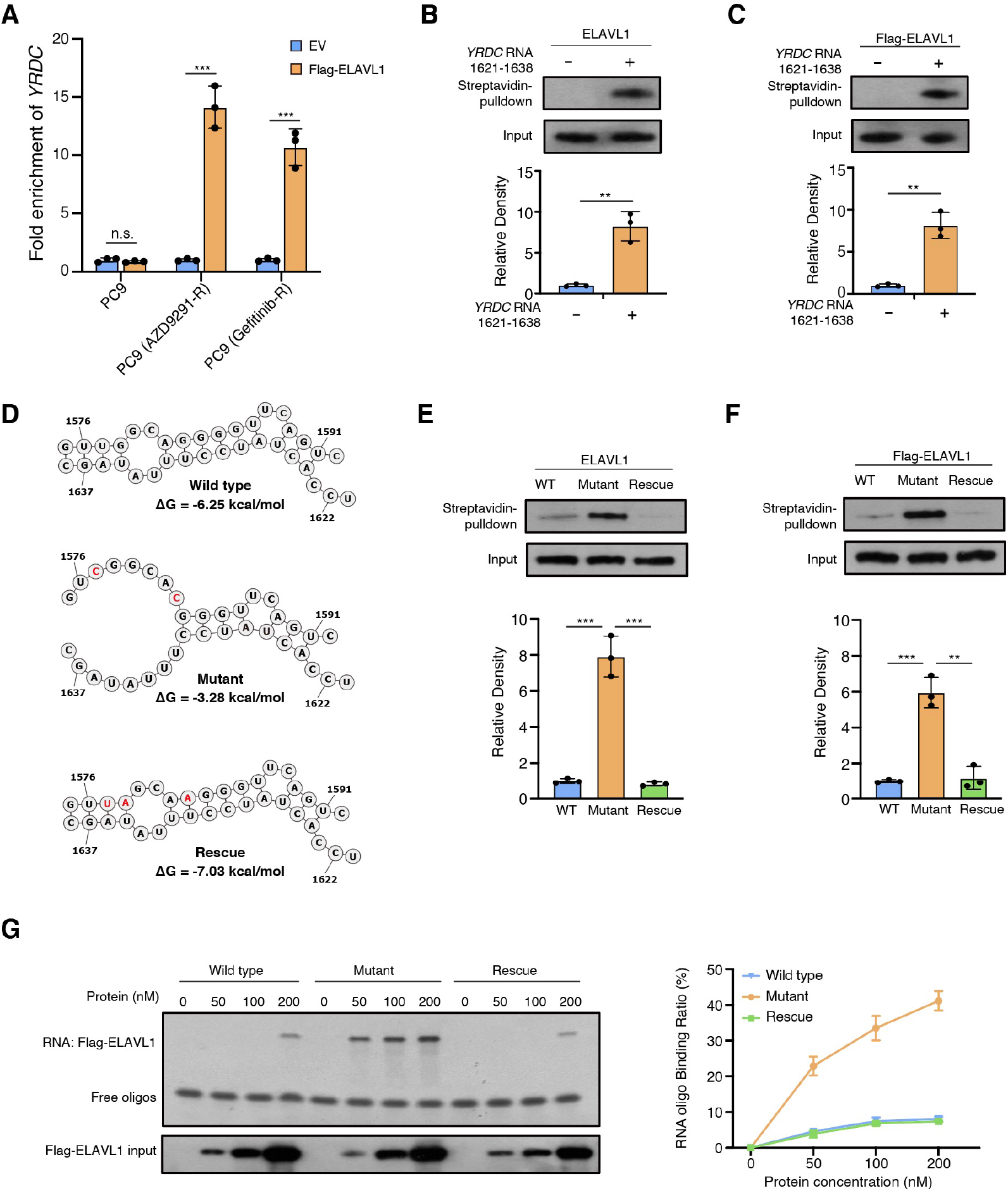
ELAVL1 Prefers to Bind Single-stranded RNA in 3’ UTR of *YRDC.* **(A)** RIP-qPCR shows the fold enrichment of ELAVL1 binding sites in 3’ UTR of *YRDC* mRNA upon Flag pull-down in PC9 cells, PC9(AZD9291-R) cells and PC9(Gefitinib-R) cells. Error bars, mean ± s.d., n = 3. *P* values were calculated using two-sided Student’s t-test, ***, *P* < 0.001, n.s., *P*, not significant. **(B)** Demonstration of endogenous ELAVL1 protein pulled down by RNA probes. Upper, western blotting; lower, quantification level. Error bars, mean ± s.d., n = 3. *P* values were calculated using Student’s *t*-test, **, *P* < 0.01. **(C)** Demonstration of purified Flag-ELAVL1 protein pulled down by RNA probes. Upper, western blotting; lower, quantification level. Error bars, mean ± s.d., n = 3. *P* values were calculated using Student’s *t*-test, **, *P* < 0.01. **(D)** The structure models of designed wild-type, mutant, and rescue RNA probes containing ARE and flanking regions. **(E)** Demonstration of endogenous ELAVL1 protein pulled down by designed endogenous RNA probes. Upper, western blotting; lower, quantification level. Error bars, mean ± s.d., n = 3. *P* values were calculated using Student’s *t*-test, **, *P* < 0.01, ***, *P* < 0.001. **(F)** Demonstration of purified Flag-ELAVL1 protein pulled down by designed endogenous RNA probes. Upper, western blotting; lower, quantification level. Error bars, mean ± s.d., n = 3. *P* values were calculated using Student’s *t*-test, **, *P* < 0.01, ***, *P* < 0.001. **(G)** EMSA (left) and line graph quantification (right) showing the binding ability of purified Flag-ELAVL1 protein with designed wild-type, mutant, and rescue RNA probes. In total, 100 nM of RNA probes was incubated with different concentrations of Flag-ELAVL1 protein. The RNA binding ratio was calculated by (RNA protein) / ((free RNA) + (RNA protein)). Error bars, mean ± s.d., n = 3.

### RNA structure switch modulates EGFR-TKIs resistance by regulating *YRDC* mRNA translation in an ELAVL1-dependent manner

For testing the sufficiency of RNA structure in ELAVL1-regulated *YRDC* mRNA translation and EGFR-TKIs resistance, *YRDC* reporter mRNAs based on CDS and 3’ UTR of *YRDC* with diverse structures (Figure 5D), the wild-type (WT) where ARE region was base-paired, the MUT disrupting ARE region’s base pairing, as well as the rescue restoring base pairing, were constructed. PC9 cells were transfected with wild-type, mutant, and rescue *YRDC* reporter plasmids under control and ELAVL1 knockdown condition. Western-blot results showed that the mutant group with a single-stranded flanking sequence was expressed higher than the wild type and rescue groups with the double-stranded flanking sequence at the protein level. In contrast, no difference was observed under the ELAVL1 knockdown condition (Figure 6A, Figure S6C). The mRNA expression of *YRDC* reporter gens was not influenced under ELAVL1 knockdown (Figure 6B), indicating ELAVL1 was not related to *YRDC* mRNA stability. The CCK8 assay also showed that the only mutant reporter genes overexpression induced a higher resistance to EGFR-TKIs, and these differences were abolished under ELAVL1 knockdown (Figures 6C, D). Thus, RNA structure modulates *YRDC* mRNA translation and EGFR-TKIs resistance via affecting ELAVL1’s accessibility to the ARE region.

**Figure 6.**
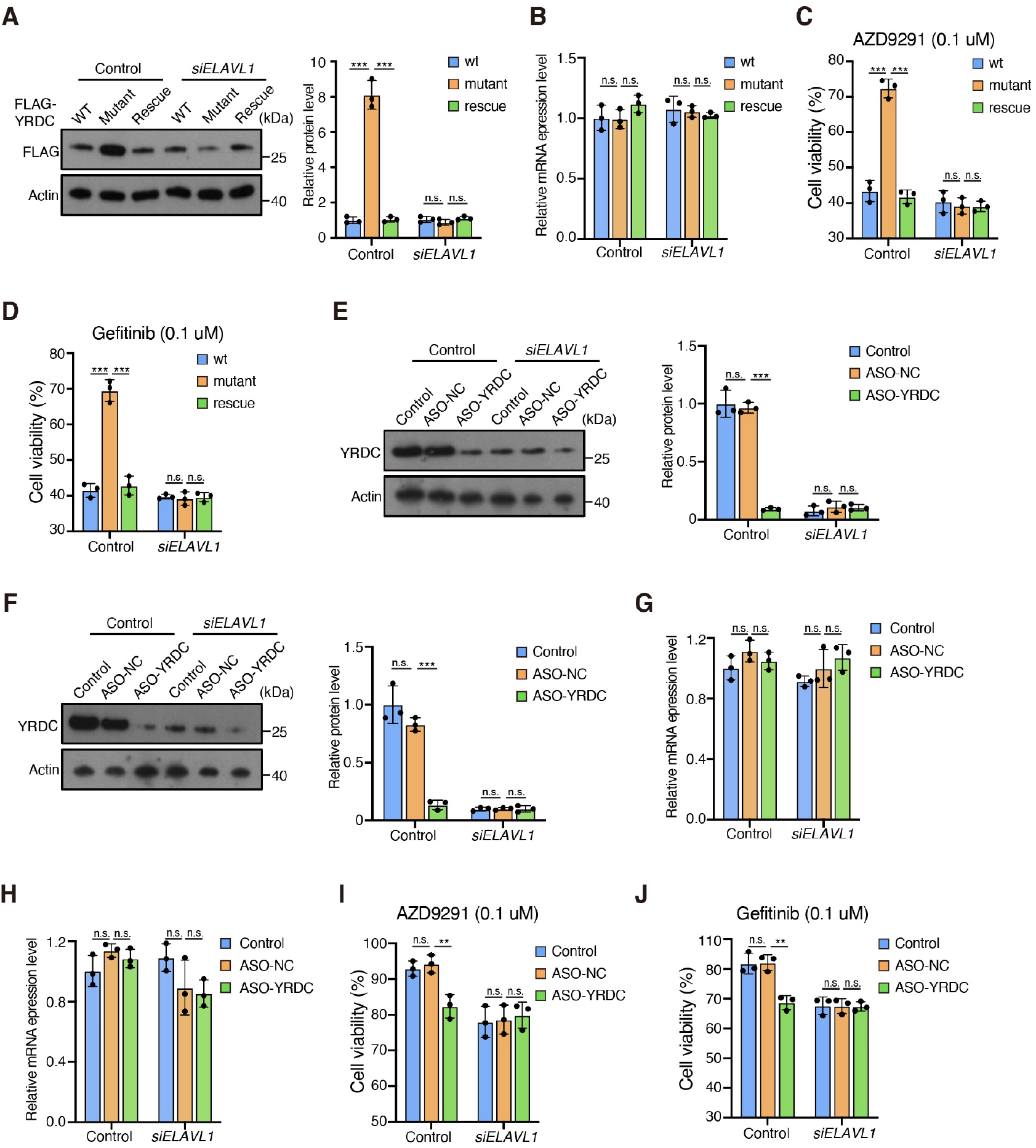
ELAVL1 Regulates *YRDC* mRNA Translation in an RNA Structure-dependent Fashion to Modulate EGFR-TKIs Resistance. **(A)** Western blot (left) and statistical (right) analyses show protein level of the wild type, mutant and rescue FLAG-YRDC upon control and ELAVL knockdown in PC9 cells. n = 3 biological replicates, error bars, mean ± SD, *P* values were calculated by the two-tailed Student’ s *t*-test, ***, *P* < 0.001, n.s., *P*, not significant. **(B)** Relative mRNA level of the wild type, mutant and rescue *FLAG-YRDC* reporter gene upon control and ELAVL knockdown in PC9 cells. n = 3, error bars, mean ± SD, *P* values were determined by the two-tailed Student’ s *t*-test, n.s., *P*, not significant. **(C,D)** CCK-8 assays for PC9 cells transfected with the wild type, mutant and rescue *FLAG-YRDC* upon control and ELAVL knockdown for 24 h followed by AZD9291 (0.1uM) (**C**) or Gefitinib (0.1uM) (**D**) treatment for another 48 h. n = 3 biological replicates, error bars, mean ± SD, *P* values were calculated by the two-tailed Student’ s *t*-test, ***, *P* < 0.001, n.s., *P*, not significant. **(E,F)** Western blot (left) and statistical (right) analyses show protein level of YRDC in PC9(AZD9291-R) (**E**) and PC9(Gefitinib-R) (**F**) cells transfected with ASO-NC or ASO-YRDC upon control and ELAVL knockdown. n = 3 biological replicates, error bars, mean ± SD, *P* values were calculated by the two-tailed Student’ s *t*-test, ***, *P* < 0.001, n.s., *P*, not significant. **(G,H)** Relative mRNA level of the *YRDC* in PC9(AZD9291-R) (**G**) and PC9(Gefitinib-R) (**H)** cells in control, ASO-NC or ASO-YRDC transfection groups upon control and ELAVL knockdown. n = 3, error bars, mean ± SD, *P* values were determined by the two-tailed Student’ s *t*-test, n.s., *P*, not significant. **(I,J)** CCK-8 assays for PC9(AZD9291-R) (**I**) and PC9(Gefitinib-R) (**J**) cells transfected with ASO-NC or ASO-YRDC upon control and ELAVL knockdown for 24 h followed by AZD9291 (0.1uM) (**I**) and Gefitinib (0.1uM) (**J**) treatment for another 48 h. n = 3 biological replicates, error bars, mean ± SD, *P* values were calculated by the two-tailed Student’ s *t*-test, ***, *P* < 0.001, n.s., *P*, not significant.

Moreover, ASO transfection was performed to test this mechanism in EGFR-TKIs resistant cells. The results showed that ASO-YRDC transfection can impair the protein expression of YRDC, and ELAVL1 knockdown abolished the difference in YRDC protein levels under ASO-NC or ASO-YRDC transfection (Figures 6E, F, Figures S6D, E). The mRNA expression level in ASO-NC or ASO-YRDC transfection groups was not influenced by ELAVL1 knockdown (Figures 6G, H). ASO-YRDC transfection also can impair the EGFR-TKIs resistance, and ELAVL1 knockdown also abolished the difference of EGFR-TKIs resistance under ASO-NC or ASO-YRDC transfection (Figures 6I, J), consistent with the YRDC protein level. Combined with our previous findings that RNA structure switch modulates ELAVL1’s accessibility to ARE region, these results suggest that ASO can be used to perturb the interaction between RNAs and RBPs, and modulating the EGFR-TKIs resistance in NSCLC cells.

To sum up, our study put forward one model that depicted the functions of RNA structural alterations in regulating *YRDC* translation and EGFR-TKIs resistance by affecting ELAVL1’s binding ability to *YRDC* (Figures 7)

**Figure 7.**
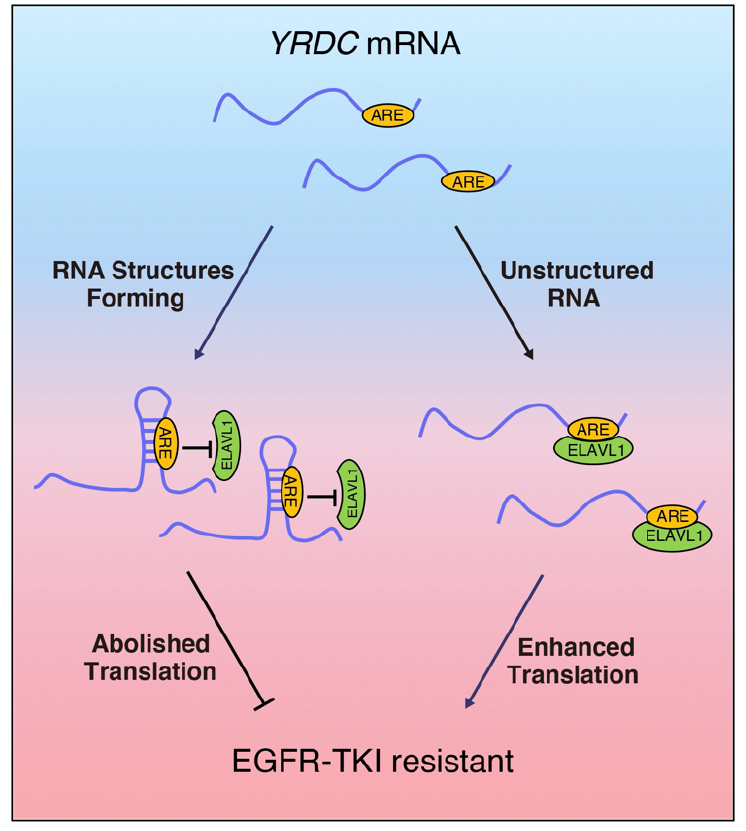
Schematic Model Shows that ELAVL1 Regulates *YRDC* mRNA Translation in a Structure-dependent Fashion to Modulate EGFR-TKIs Resistance.

## DISCUSSION

In this study, transcriptome-scale RNA structures were profiled using an acquired resistant model of NSCLC cells that reveal an RNA structure switching-dependent mechanism in regulating EGFR-TKI resistance. Intriguingly, we show that (1) RNA structural regions are enriched in UTR, translation start site, and stop codon, indicating its role of RNA post-transcriptional regulation and translation control, (2) by combining with Ribosome profiling data, we found that RNA translation efficiencies are dynamic during the period of drug resistance, and impaired by the RNA structures, (3) the RNA structure within *YRDC* mRNA’s 3’ UTR could modulate NSCLC cell translation and EGFR-TKIs resistance by regulating ELAVL1 binding. Overall, our study demonstrates RNA structure-based regulation of translation control, critical for the drug sensitivity to resistance in NSCLC cells. Furthermore, RNA structures seem to be a molecular switch for controlling translation in cancer drug resistance, thereby allowing us to overcome drug resistance by modulating RNA structure.

LC accounts for a major factor resulting in cancer-associated death globally, while NSCLC is a frequent subtype(Siegel et al., 2021). Nearly 2/3 of NSCLC cases harbor the carcinogenic driver mutation(Rotow and Bivona, 2017), and the TKIs for sensitizing EGFR mutations are investigated widely. Nevertheless, many cases acquired EGFR-TKIs resistance in a short time. The mechanisms of resistance are divided into ‘on-target’ and ‘off-target’(McGranahan and Swanton, 2017). The former occurs in the case of a changed drug’s primary target, which limits the target inhibition capacity of the drug, among which the T790M point mutation of EGFR is the most prevalent for gefitinib, erlotinib, or afatinib resistance. Third-generation EGFR-TKI (AZD9291) can solve the problem of resistance to EGFR T790M mutation; however, acquired resistance is inevitable(Sullivan and Planchard, 2016). The accumulating evidence indicated that the On-target resistance was insufficient to explain the resistance of EGFR-TKIs. By contrast, off-target resistance occurs by activating collateral signaling in the downstream signaling or parallel to the signaling via a driver oncoprotein. Therefore, exploring the new mechanism for overcoming the resistance to EGFR-TKIs in LC is crucial for therapy. Our work profiled RNA structures atlas during EGFR-TKIs resistance in NSCLC cells, which provided a new perspective that RNA structural switch-mediated translation control modulates drug resistance. Meanwhile, this study used gefitinib and AZD9291 (the first- and third-generation EGFR TKIs, separately) for drug resistant-cells construction to obtain the comprehensive RNA structure atlases and general regulation mechanism in EGFR-TKIs resistance.

RNA structures are involved in almost all RNA processing, such as polyadenylation, splicing, localization, degradation, and translation(Wang et al., 2021b). Moreover, recent studies proved that RNA structures are crucial in physiological processes, including embryogenesis(Beaudoin et al., 2018; Shi et al., 2022; Shi et al., 2020), cardiac specification(Xue et al., 2016), neurogenesis(Wang et al., 2021a), viral infection(Cao et al., 2021; Huston et al., 2021; Mizrahi et al., 2018; Sun et al., 2021; Ziv et al., 2020). Although RNA structures are related to RNA processing and essential for RNA functions, gene regulation based on RNA structure in cancer drug resistance remains unknown. This work constructed an RNA structure atlas in NSCLC cells, associating RNA structure with post-transcriptional regulation. Firstly, RNA structures were found enriched in UTRs. Previous studies have reported that conserved structured elements found in 35% of UTRs(Mustoe et al., 2018), indicating that the RNA structures in UTRs are crucial for RNA fate determination and function. Secondly, RNA structures were found to impair translation efficiency in NSCLC cells. RNAs that had more structure within 5’ UTR decreased the binding affinity of ribosomes, lowering translation efficiency(Mustoe et al., 2018). Meanwhile, ribosomes have an essential effect on unlocking secondary structures in CDS(Beaudoin et al., 2018; Mustoe et al., 2018; Shi et al., 2020). Additionally, RNAs with a double-stranded structure within 3’ UTR showed a lower translation efficiency, possibly due to their higher accessibility to RNA decomposition and translation inhibition mechanisms(Wang et al., 2021b).

*YRDC* is involved primarily in adenosine 37 *N^6^*-threonyl-carbamoylation in tRNA’s ANN-type tRNAs (t^6^A) synthesis for recognizing ANN codons(El Yacoubi et al., 2009). t^6^A represents the widely distributed modification necessary to maintain the accurate and efficient translation(Lescrinier et al., 2006; Murphy et al., 2004), and *YRDC* regulates HCC cell resistance to lenvatinib by regulating KRAS translation(Guo et al., 2021). The present work found that *YRDC* facilitates EGFR-TKIs resistance in NSCLC cells, and the RNA structure in 3’ UTR of *YRDC* mRNA can modulate translation efficiency, which only exists in PC9 cell that is sensitive to EGFR-TKIs. Further research found that this structure inhibits the binding affinity of ELAVL1 and impairs the *YRDC* translation. ELAVL1 is crucial for mRNA stability and translation control(Hinman and Lou, 2008) by binding the AU- or U-rich sequences in the corresponding 3’ and 5’ UTRs(López de Silanes et al., 2004). Many studies revealed the involvement of ELAVL1 in post-transcriptional regulation of several specific cancers(Abdelmohsen and Gorospe, 2010; Schultz et al., 2020). Moreover, the RNA structure greatly affects the binding affinity of its zebrafish ortholog Elavl1a to the AUrich motif and is essential for zebrafish embryogenesis(Shi et al., 2020). Our study performed RNA pulldown assays *in-vivo* and *in-vitro* using structural mutant and rescue form of 3’ UTR of *YRDC.* The results showed that ELAVL1 protein preferentially binds to single-strand RNA of 3’ UTR of *YRDC*, concordant with the result of the EMSA assay. Moreover, we found that the RNA structural switch in 3’ UTR of *YRDC* modulates the ELAVL1 binding, essential for controlling *YRDC* translation.

Previous studies reported several mechanisms that regulate translational modulation for shaping tumor development and treatment resistance(Fabbri et al., 2021). The translation modulation by RNA structure reconstruction can be exploited in cancer treatment. Due to the progression of medicinal chemistry, targeted delivery, and molecular mechanisms, ASOs have substantially improved efficacy and properties(Crooke et al., 2021). Besides inducing target RNA degradation, ASO can also modulate splicing, translation, polyadenylation, etc(Crooke et al., 2021; Roberts et al., 2020). A recent study showed that ASO could disturb the RBP-RNA target interactions by reconstructing RNA structure(Sun et al., 2021). In our study, we found the ASO transfection could decrease the *YRDC* translation and resistance to EGFR-TKIs, indicating that the ASO-mediated RNA structural switch serves as a possible way for overcoming drug resistance in cancer.

Although this study provides a comprehensive resource of RNA structures in EGFR-TKI resistance model, and uses this information to explore the potential RNA structure-dependent mechanism in drug resistance, there are nevertheless a few limitations, stemming both from the samples for RNA structure detection and regarding the validations of the ASO treatment. First, the RNA structural information obtained from the cellular models with EGFR-TKI resistance and sensitivity that were constructed by using a representative NSCLC cell line, PC9. The LC was highly heterogeneous between different people, although we used two EGFR-TKI resistant cell line to profile RNA structures, it was difficult to represent the RNA structures in all LC cells. Thus, detection the RNA structures in different cell lines or cancer samples could help in elucidating exactly how dynamic RNA structures contributes to EGFR-TKI resistance. Second, although we have demonstrated that the RNA structure-dependent mechanism in drug resistance and potential ASO treatment in different cells, their mechanisms and the action of ASO should be studied further, and their efficacy and side-effects must be assessed by in vivo validations using animal models.

Collectively, our study profiles the RNA structure landscape in cells with EGFR-TKI resistance and sensitivity, and reveals the novel RNA structure switching-dependent mechanism in regulating EGFR-TKI resistance. Notably, the usage of ASO in perturbing the interaction between RNA and protein provide a potential strategy for cancer therapy. Thus, we reasonably speculate that the RNA structure-dependent mechanism and RNA structure targeted drugs could be used for the clinical treatment of NSCLC.

## STAR METHODS

### KEY RESOURCES TABLE

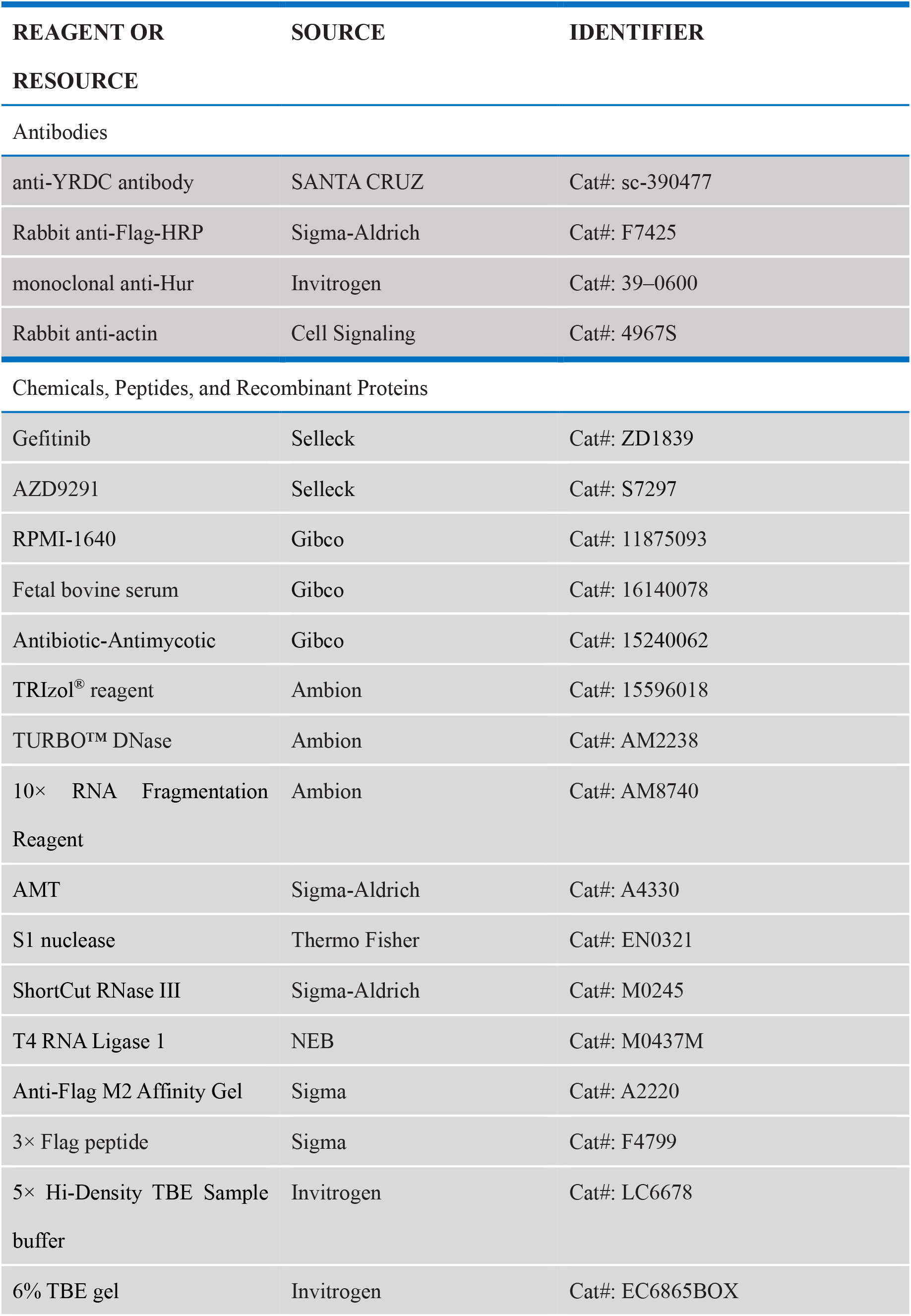
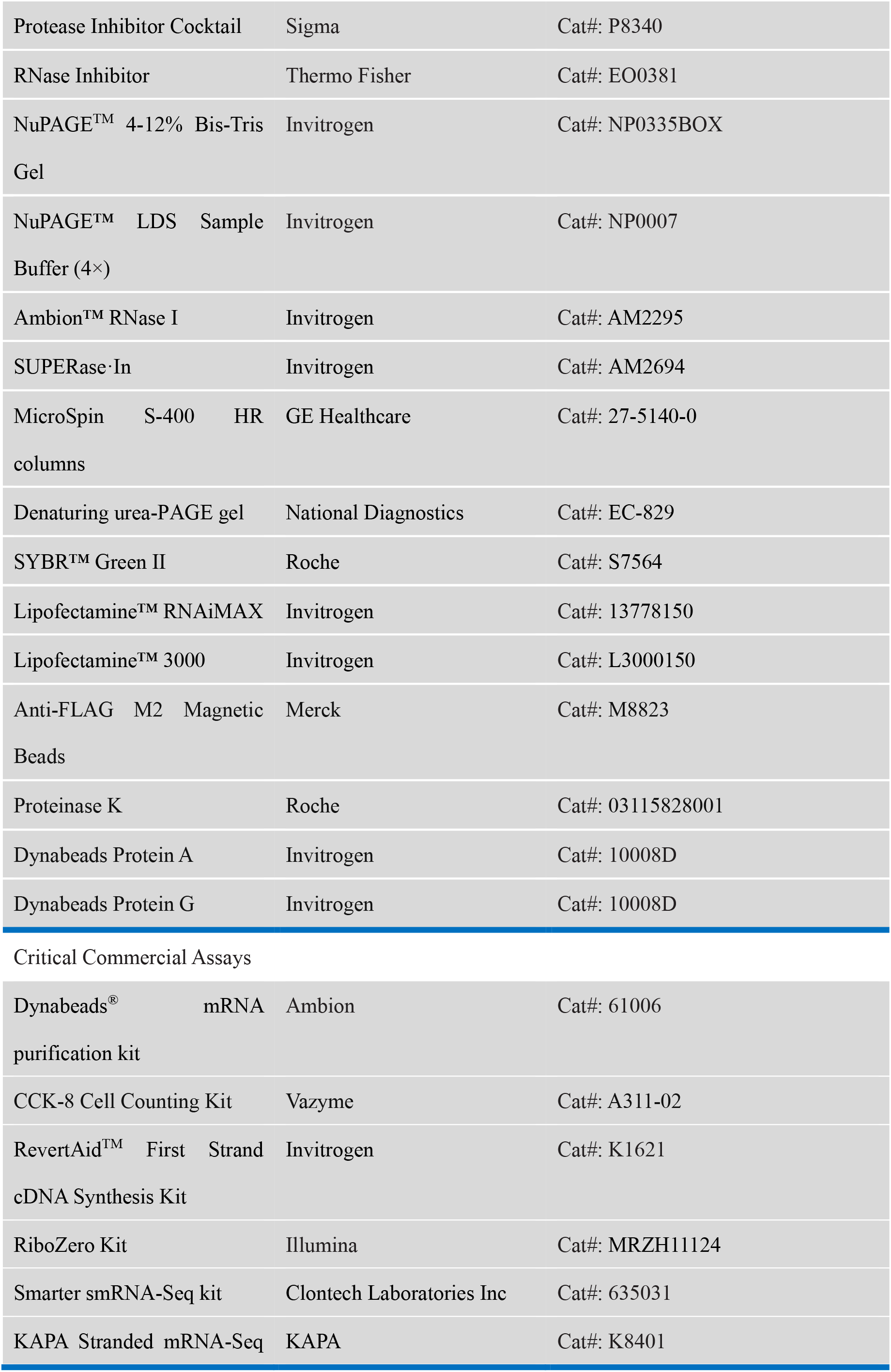
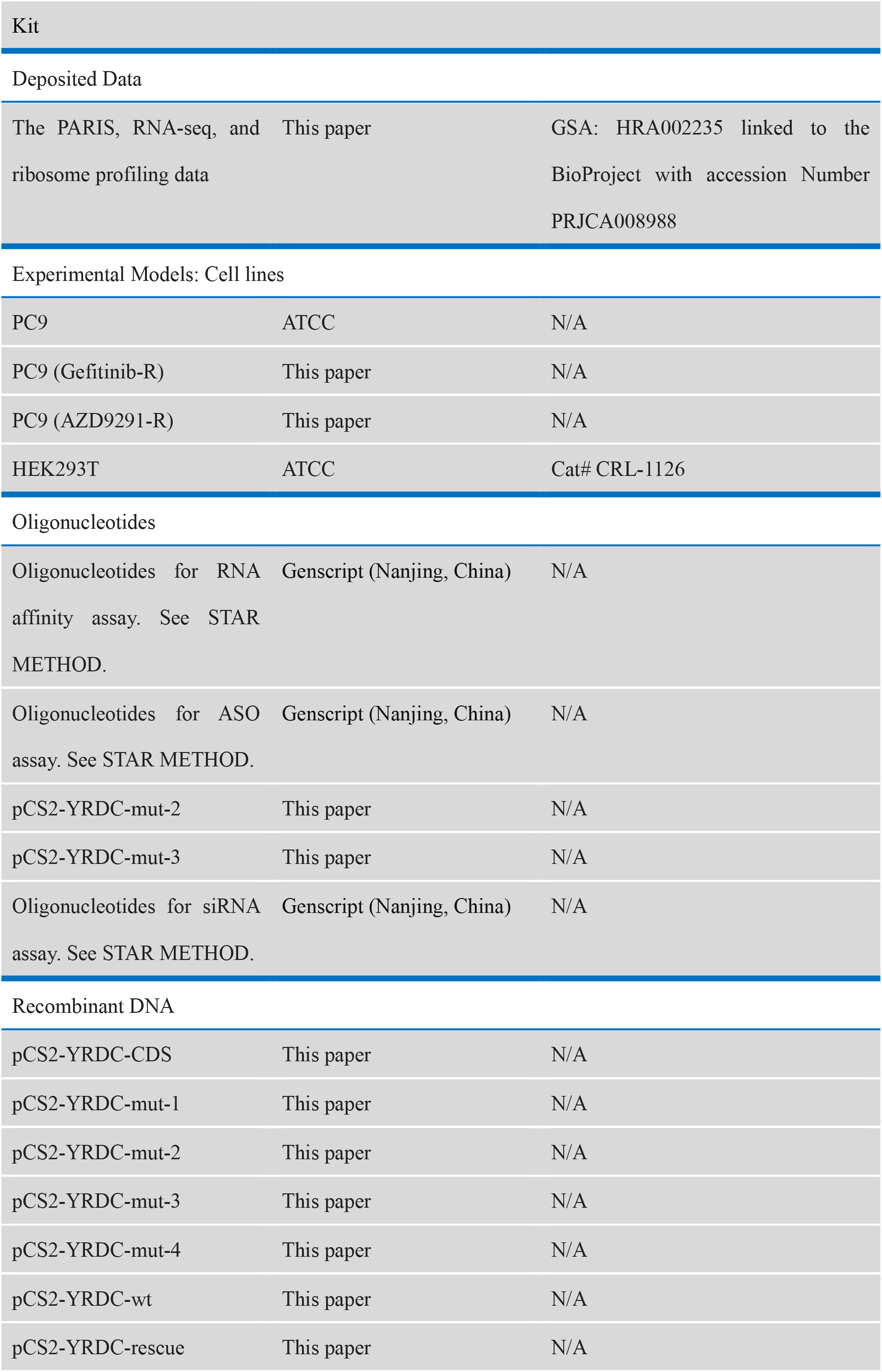
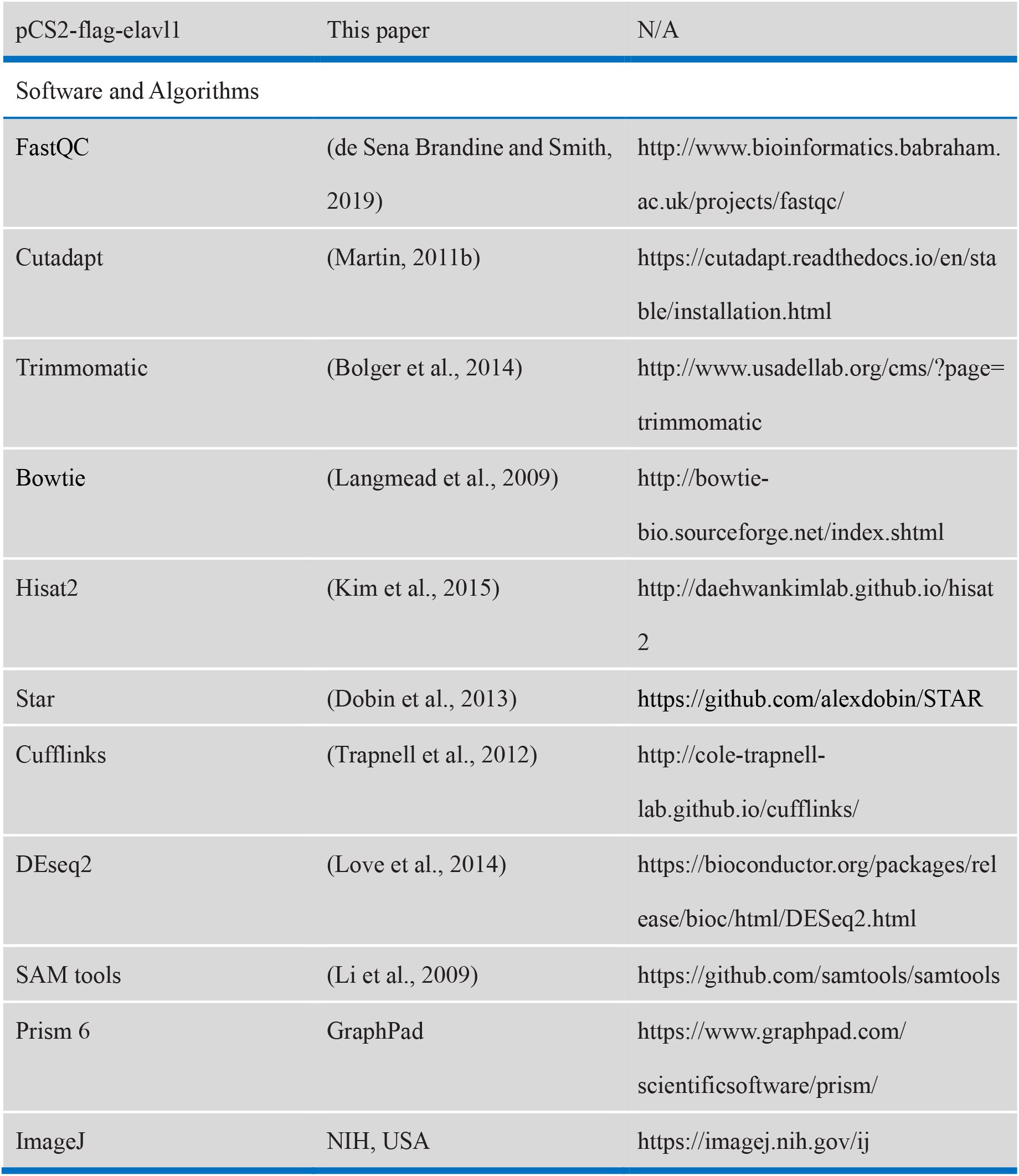

### RESOURCE AVAILABILITY

#### Lead Contact

Further information and requests for resources and reagents should be directed to and will be fulfilled by the Lead Contact, Quancheng Kan

#### Materials Availability

All unique/stable reagents generated in this study are available from the Lead Contact with a completed Materials Transfer Agreement.

#### Data and Code Availability

The PARIS, RNA-seq, and ribosome profiling data supporting the conclusions of this article have been deposited in the Genome Sequence Archive (GSA: HRA002235 linked to the BioProject with accession Number PRJCA008988). The code is available on GitHub (https://github.com/ankecode/drug_resistance).

### EXPERIMENTAL MODEL AND SUBJECT DETAILS

#### Cell lines and culture conditions

NSCLC cell lines PC-9 was newly purchased from American Type Culture Collection. PC9 and HCC827 cells harbored EGFR exon 19 deletion. PC9 (Gefitinib-R) and PC9 (AZD9291-R) were passaged with low concentration of Gefitinib (Selleck, ZD1839) or AZD9291 (Selleck, S7297) (0.1 μM) and sequentially cultured in increasing concentrations of these TKIs (0.3, 1 μM). Cell lines were grown in RPMI-1640 (Gibco, 11875093) supplemented with 10% fetal bovine serum (FBS, Gibco, 16140078) and Antibiotic-Antimycotic (Gibco by Life Technologies, 15240062) at 37 °C under 5% CO2.

### METHOD DETAILS

#### PARIS Experimental Method

PARIS experiments were performed as previously reported(Lu et al., 2016). PC9, PC9 (Gefitinib-R) and PC9 (AZD9291-R) cells were treated with or without AMT and crosslinked with 365 nm UV. Cell lysates were digested with S1 nuclease and RNA purified using TRIzol. Purified RNA was further digested with ShortCut RNase III to smaller fragments. RNA was separated by 12% native polyacrylamide gel and then the first-dimension gel slices were further electrophoresed in a second dimension 20% urea-denatured gel. Crosslinked RNA above the main diagonal was eluted, proximity ligated with T4 RNA ligase I and photo-reversed with 254 nm UV. Ribosomal RNA fragments were removed using the RiboZero Kit (Illumina, MRZH11124). The proximity-ligated RNA molecules were subjected to library preparation using the Smarter smRNA-Seq kit (Clontech Laboratories Inc).

#### Western blot and immunofluorescence

Western blot was performed as previously reported(Heng et al., 2020) using the following antibodies: anti-YRDC antibody (SANTA CRUZ, sc-390477), monoclonal anti-Hur (Invitrogen, 39–0600), anti-β-actin antibody (Cell Signaling Technology, 4967, RRID:AB_330288), anti-Flag antibody (Sigma-Aldrich, F7425).

#### Protein purification in mammalian cells

HEK293 cells were transfected with Flag-tagged *ELAVL1* plasmids using the PEI transfection reagent (MKbio). After 48 h, cells were lysed with lysis buffer (50 mM Tris-HCl, pH 7.4, 150 mM NaCl, 1% NP-40, Protease inhibitor cocktail) and sonicated (10% output, 10 s pulse-on, 20 s pulse-off) for 1 min by a Sonic Dismembrator (Thermo Fisher). After removing cell debris through centrifugation at 13,300 rpm for 20 min, the lysates were incubated with the anti-Flag M2 Affinity Gel (Sigma-Aldrich, A2220) for 4 h, at 4°C. Afterwards the samples were washed five times with lysis buffer and twice with TBS buffer (20 mM Tris-HCl pH 7.4, 150 mM NaCl). The gel-bound proteins were eluted with a 4 mg/ml 3× Flag peptide (Sigma-Aldrich, F4799) by incubating the mixture for 1 h, at 4°C. The eluate containing purified protein was concentrated using VIVASPIN (Sartorius Stadium Biotech) and quantified by Coomassie brilliant blue staining and Western blot analysis.

#### RNA-seq

Total RNA was isolated from cells with TRIzol reagent and mRNA was purified using the Dynabeads mRNA purification kit (Ambion, 61006). Fragmented mRNA was used for library construction using the KAPA Stranded mRNA-Seq Kit (KAPA, K8401), according to the manufacturer’s protocol.

#### Ribosome profiling experiments

Ribosome profiling experiments were performed as previously reported(Calviello et al., 2016), with some modifications. Cells were incubated in 100 mg/ml medium for 5 min at 37 °C. Cells were then transferred into 200 μl of ice-cold lysis buffer. After triturating the cells ten times through a 26-G needle, the lysate was clarified by centrifugation for 10 min at 20,000 g at 4°C to recover the soluble supernatant. 1 μl of RNase I (100 U/μl) was added to 200 μl of lysate. After a 45-min incubation at room temperature with gentle mixing, 10 μl of SUPERase·In (Invitrogen, AM2694) RNase inhibitor was added to stop the nuclease digestion. MicroSpin S-400 HR columns (GE Healthcare, 27-5140-01) were equilibrated with 3 ml of mammalian polysome buffer by gravity flow and emptied by centrifugation at 600 g for 4 min. Immediately 100 μl of the digested lysate was loaded on the column and eluted by centrifugation at 600 g for 2 min. RNA was extracted from the flow-through (approximately 125 μl) using Trizol LS (Life Technologies, 10296-010). Ribosomal RNA fragments were removed using the RiboZero Kit (Illumina, MRZH11124) and were separated on a 17% denaturing urea-PAGE gel (National Diagnostics, EC-829). RNA fragments sized 27 nt to 30 nt were cut out of the gel, and were subjected to library preparation using the Smarter smRNA-Seq kit (Clontech Laboratories Inc).

#### Reverse-transcription quantitative PCR

Reverse-transcription quantitative PCR (RT-qPCR) was carried out to examine the relative abundance of target RNA. 0.1 μg of total RNA was used for cDNA synthesis using the RevertAid™ First Strand cDNA Synthesis Kit (Thermo). Experiments were performed with the Takara SYBR Premix Ex Taq (Takara) according to the manufacturer’s instructions and examined by a CFX96 Real-Time PCR System (Bio-Rad). The primers used for RT-qPCR in this study are listed as follows:

YRDC-F: GCCTCTTGTAGGCATTCGGA
YRDC-R: AGTACTTTCCAGGGCACAGC

#### In vivo RNA Pulldown Assay

The biotin-labeled RNA probes were synthesized by the GenScript Biotech Corp. In vivo RNA pull-down assays were carried out using cells extracts as previously described (Yang et al., 2017) with some modifications. RNA was heated in metal-free water for 2 min at 95 °C. The RNA was then flash-cooled on ice. The RNA 3X SHAPE buffer (333 mM HEPES, pH 8.0, 20 mM MgCl2, 333 mM NaCl) was added and the RNA was allowed to equilibrate at 37 °C for 10 min. Cell extracts were precleared for 1 h at 4 °C by incubation with streptavidin-conjugated magnetic beads (NEB) in binding buffer (50 mM Tris-HCl pH 7.5, 250 mM NaCl, 0.4 mM EDTA, 0.1% NP-40, 1 mM DTT) supplemented with 0.4 U/μl RNasin (Promega). Biotin-labeled RNA oligonucleotides were incubated with pre-cleared nuclear extracts for 2 h at 4 °C under gentle rotation together with streptavidin-conjugated magnetic beads which were pre-cleared by incubation with 0.2 mg/ml tRNA (Sigma) and 0.2 mg/ml BSA (Amresco) for 1 h at 4 °C under gentle rotation. Beads were washed three times with wash buffer (50 mM Tris-HCl pH 7.5, 250 mM NaCl, 0.4 mM EDTA, 0.1% NP-40, 1 mM DTT, 0.4 U/μl RNasin (Promega)). For western blotting analysis, samples were separated on SDS-PAGE and transferred onto PVDF membrane. After blocking with 5% non-fat milk in TBST for 1 h, the membrane was then incubated for 1 h at 4 °C with Monoclonal anti-Hur (Invitrogen, 39-0600) diluted at 1:1 000 in 5% milk. Protein levels were visualized using ECL Western Blotting Detection Kit (GE Healthcare)

#### In vitro RNA Pulldown Assay

In vitro RNA pulldown assay was performed according to the previously reported method (Yang et al., 2017) with some modifications. Generally, RNA was heated in metal-free water for 2 min at 95 °C. The RNA was then flash-cooled on ice. The RNA 3X SHAPE buffer (333 mM HEPES, pH 8.0, 20 mM MgCl2, 333 mM NaCl) was added and the RNA was allowed to equilibrate at 37 °C for 10 min. Then, 10 pmol of purified Flag-ELAVL1 protein and 10 pmol of biotin-labeled probes were incubated with 15 μl streptavidin-conjugated magnetic beads (NEB) in binding buffer (50 mM Tris-HCl pH 7.5, 250 mM NaCl, 0.4 mM EDTA, 0.1% NP-40, 1 mM DTT, 0.4 U/μl RNase inhibitor) for 1 h at 4 °C. After washing with binding buffer for three times, the beads-bound proteins were heated in NuPAGE™ LDS Sample Buffer (4 ×) (Invitrogen) and then separated on the NuPAGE™ 4%-12% Bis-Tris Gel (Invitrogen), and subjected to western blotting analysis with anti-Flag antibody (Sigma-Aldrich; RRID: AB_439687)

#### Electrophoretic Mobility Shift Assay (EMSA)

Purified Flag-tagged ELAVL1 proteins were diluted to a series of concentrations of 0.2 μM, 0.5 μM, 1 μM, and 2 μM in binding buffer (50 mM Tris-HCl pH 7.5, 100 mM NaCl, 0.4 mM EDTA, 0.1% NP-40, and 40 U/ml RNasin, 1 mM DTT, 50% glycerol, 5 ng/μl BSA). 1 μl synthesized Cy3-labeled RNA probes (100 nM final concentration) and 1 μl purified protein (10 nM, 50 nM, 100 nM, and 200 nM final concentration, respectively) were mixed and incubated at room temperature for 30 min. Then, 1 μl glutaraldehyde (0.2% final concentration) was added into the mixture which was incubated at room temperature for 15 min. The entire 11 μl RNA-protein mixture was mixed with 5 μl 5× Hi-Density TBE Sample buffer and separated on 6% TBE gel on ice for 30 min at 80 V. The gel was scanned on a Typhoon 9400 (GE Healthcare, USA) imager. Quantification of each band was carried out using ImageJ. The RNA binding ratio at each protein concentration was determined by (RNA-protein)/((free RNA) + (RNA-protein)).

#### siRNA and ASO transfection

si*ELAVL1*: 5’-AAGAGGCAAUUACCAGUUUCA-3’, si*YRDC*: 5’-CAUUCGGAUUCCUGAUCAU-3’, siNC: 5’-UUCUCCGAACGUGUCACGU-3’, ASO-*YRDC*: 5’-/i2MOErC/*/i2MOErC/*/i2MOErC/*/i2MOErT//i2MOErG//i2MOErG//i2MOErC//i2 MOErT//i2MOErA//i2MOErT//i2MOErA//i2MOErA//i2MOErA//i2MOErG//i2MOE rG//i2MOErA//i2MOErT/*/i2MOErA/-3’, ASO-NC: 5’-/i2MOErA/*/i2MOErT/*/i2MOErG/*/i2MOErT//i2MOErG//i2MOErT//i2MOErC//i2 MOErC//i2MOErT//i2MOErG//i2MOErT//i2MOErT//i2MOErA//i2MOErA//i2MOEr C//i2MOErT//i2MOErC//i2MOErA/*/i2MOErT/*/i2MOErC/*/i2MOErA/-3’ were synthesized by Genscript (Nanjing, China). siRNAs and ASOs were transfected by using Lipofectamine RNAiMAX (Life technologies), and used for drug treatment after 24h.

#### CCK8 assay

Cells grown in 96-well plates were incubated with various drug concentrations for 48 hours. Cell viability was measured by a CCK-8 Cell Counting Kit (Vazyme, A311-02) and calculated using the GraphPad Prism software.

#### ELAVL1 RIP-qPCR

PC9, PC9 (Gefitinib-R) and PC9 (AZD9291-R) cells were transfected with Flag-tagged ELAVL1 plasmids using the Lipofectamine™ 3000 (Life technologies). And cells were harvested after 48h and lysed in NETN lysis buffer (150mM NaCl, 0.5% NP-40, 50mM Tris-HCl, pH 7.4). In brief, lysate was incubated with Anti-FLAG M2 Magnetic Beads (Merck, M8823) for 4 h at 4 °C. After washing, proteins were digested by 4 μg μl^-1^ proteinase K (Roche, 03115828001) in 200μl PK buffer (50mM NaCl, 100mM Tris-HCl pH7.4, 10mM EDTA) for 20 min at 37 °C, followed by incubation with 200 μl PK-urea buffer (100 mM Tris-HCl pH 7.4, 50mM NaCl, 10mM EDTA, 7M urea) for 20min at 37°C. After washing, RNA was collected by EtOH precipitation and then used for RT-qPCR.

#### RNA-seq data processing and analysis

The raw sequencing reads were processed using FastQC (http://www.bioinformatics.babraham.ac.uk/projects/fastqc/). Low-quality reads and shorter reads were trimmed by cutadapt (V 3.0) and Trimmomatic (V 0.39).(Bolger et al., 2014) Processed reads were mapped to the human genome (GRCh38) from the Ensemble annotation using Hisat2 (V 2.2.1) with “-p -N –dta”. After quality filtering (≥ 20) using SAM tools (V1.11),(Li et al., 2009) read counts and corresponding RPKMs were calculated. The fold change was calculated using the DEseq2 (V 1.26.0) package.

#### Ribosome profiling RNA-seq data processing and analysis

Reads sequencing data were used to obtain ribosome protected fragments RNA-seq data. The quality of raw sequencing reads was evaluated using FastQC (http://www.bioinformatics.babraham.ac.uk/projects/fastqc/). Low-quality reads were t filtered by cutadapt (V 3.0)(Martin, 2011a) and Trimmomatic (V 0.39). Filtered reads were mapped to the human rRNA transcriptome using Bowtie (v 1.3.0)(Langmead et al., 2009) and unmapped reads were retained. The retained reads were mapped to human genome (GRCh38) using Bowtie (v 1.3.0). The mRNAs expression levels were calculated by Cufflinks (v2.2.1) with ‘‘-p -u -G”. Translation efficiency of each mRNA was calculated by the ratio of RPFs and the mean expression of input mRNA replicates. Translation efficiency (TE) was calculated by (RPKM of Ribo-seq)/(RPKM of RNA-seq).

#### PARIS data processing and analysis

After the sequencing of PARIS libraries, raw sequencing reads were evaluated using FastQC (http://www.bioinformatics.babraham.ac.uk/projects/fastqc/). The low quality and shorter reads were trimmed by cutadapt (V 3.0) and Trimmomatic (V 0.39). Processed reads were mapped to the human genome (GRCh38) from the Ensembl annotation using Star (V 2.5.3) with “--outReadsUnmapped Fastx -- outFilterMultimapNmax 10 --outSAMattributes All --alignIntronMin 1 -- outSAMmultNmax 2 --chimOutType WithinBAM SoftClip --outSAMtype BAM Unsorted --scoreGapNoncan -4 --scoreGapATAC -4 --chimSegmentMin 15 -- limitOutSJcollapsed 9000000 --limitIObufferSize 950000000 -- chimJunctionOverhangMin 15 --runThreadN 32 “. The sam file was converted to bam file using SAM tools (V1.11). The gap regions and duplex group were classified by icSHAPE-pipe (https://github.com/lipan6461188/icSHAPE-pipe).

### QUANTIFICATION AND STATISTICAL ANALYSIS

Unless otherwise indicated, data are presented as mean ± SD of 3 independent experiments. All statistical analyses were performed with Graph Prism 6.0 software, and the statistics were analyzed by unpaired Student’s t test. The correlation between genes expression was analyzed using Pearson’s test. *p* values were provided as *\*P* < 0.05, \*\**P* < 0.01, \*\*\**P* < 0.001.

## Data availability

The PARIS, RNA-seq, and ribosome profiling data supporting the conclusions of this article have been deposited in the Genome Sequence Archive (GSA: HRA002235 linked to the BioProject with accession Number PRJCA008988).

## Code availability

The code is available on GitHub (https://github.com/ankecode/drug_resistance).

## SUPPLEMENTAL INFORMATION

Figure S1-S6

Table S1 Summary and statistics of PARIS, ribosome profiling and RNA-seq data

Table S2 GO term enrichment analysis of common changed genes compared between EGFR-TKI resistant cells and EGFR-TKI sensitive cells.

## ACKNOWLEDGEMENTS

This work was supported by grants from the Province and Ministry Coconstruction Major Program of Medical Science and Technique Foundation of Henan Province (No. SBGJ202001007), the National Natural Science Foundation of China (31870809, 32121001), and the Special Fund for Young and Middle School Leaders of Henan Health Commission (HNSWJW-2020017).

## AUTHOR CONTRIBUTIONS

Q.K., X.T. and Y.-G.Y. conceived this project, supervised the study; B.S. performed the experiments; Y.W. constructed drug resistant cell lines; K. A. and B..S. performed bioinformatics analysis with assistance from Y.F. and Q.C.Z.; B.S. wrote the manuscript with input from all the other authors. All authors read and approved the final manuscript.

## Supplemental Information

**Figure S1.**
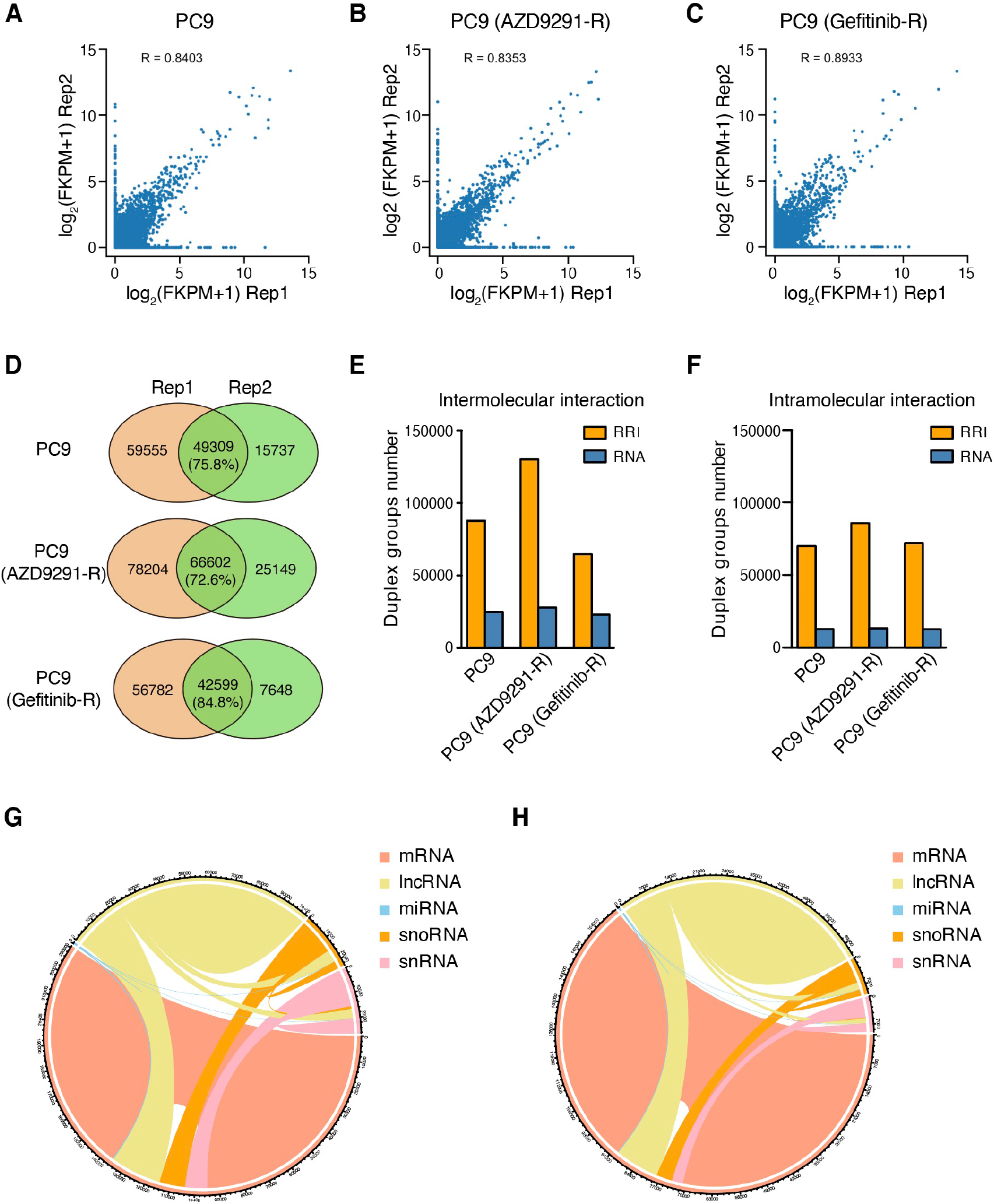
Validation of PARIS data quality, related to Figure 1. **(A-C)** RPKM correlation of PARIS data between biological replicates for PC9 cells (**A**), PC9(AZD9291-R) cells (**B**) and PC9(Gefitinib-R) cells (**C**). **(D)** Overlay of RNA duplex groups between biological replicates. **(E)** The number of intermolecular RNA-RNA duplexes and transcripts in EGFR-TKIs resistant cells and sensitive cells. **(F)** The number of intramolecular RNA-RNA duplexes and transcripts in EGFR-TKIs resistant cells and sensitive cells. **(G,H)** Circos plot of the landscape of RNA-RNA duplexes detected by PARIS in PC9(AZD9291-R) cells (**G**) and PC9(Gefitinib-R) cells (**H**).

**Figure S2.**
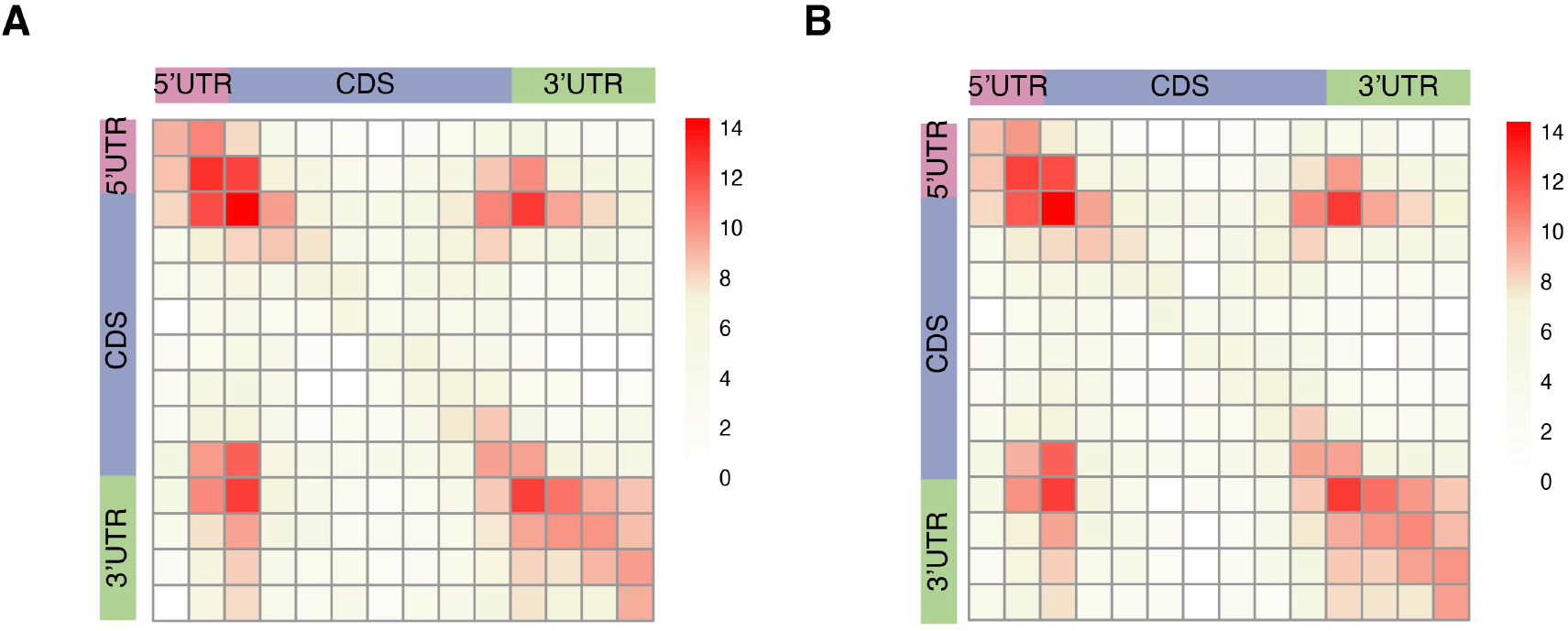
RNA structure regions are enriched in UTR, translation initiation sites, and stop codons, related to Figure 2. **(A,B)** Two-dimensional heatmap showing enrichment of mRNA structures based on the location of chimera ends in PC9(AZD9291-R) cells (**A**) and PC9(Gefitinib-R) (**B**) cells

**Figure S3.**
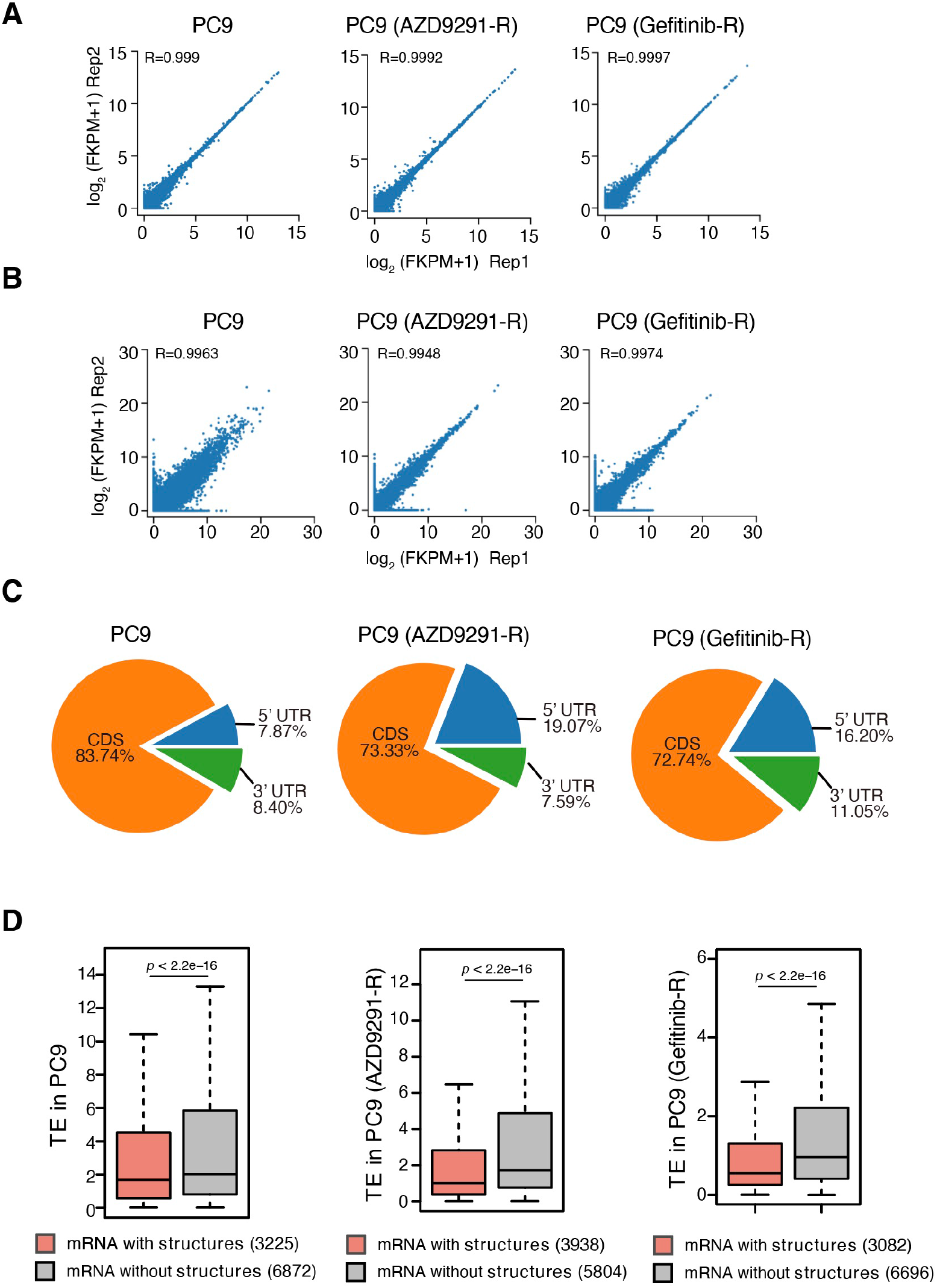
RNA structures correlate with RNA translation efficiency, related to Figure 2. **(A)** RPKM correlation of RNA-seq data between biological replicates. **(B)** RPKM correlation of Ribo-seq data between biological replicates. **(C)** Pie chart illustrating the proportion of Ribosome profiling sequencing data in three transcripts regions. **(D)** Boxplot chart showing decreased translation efficiency (TE) for mRNAs displaying RNA structures compared to mRNAs without RNA structures. *P* values were calculated by the Wilcox. test.

**Figure S4.**
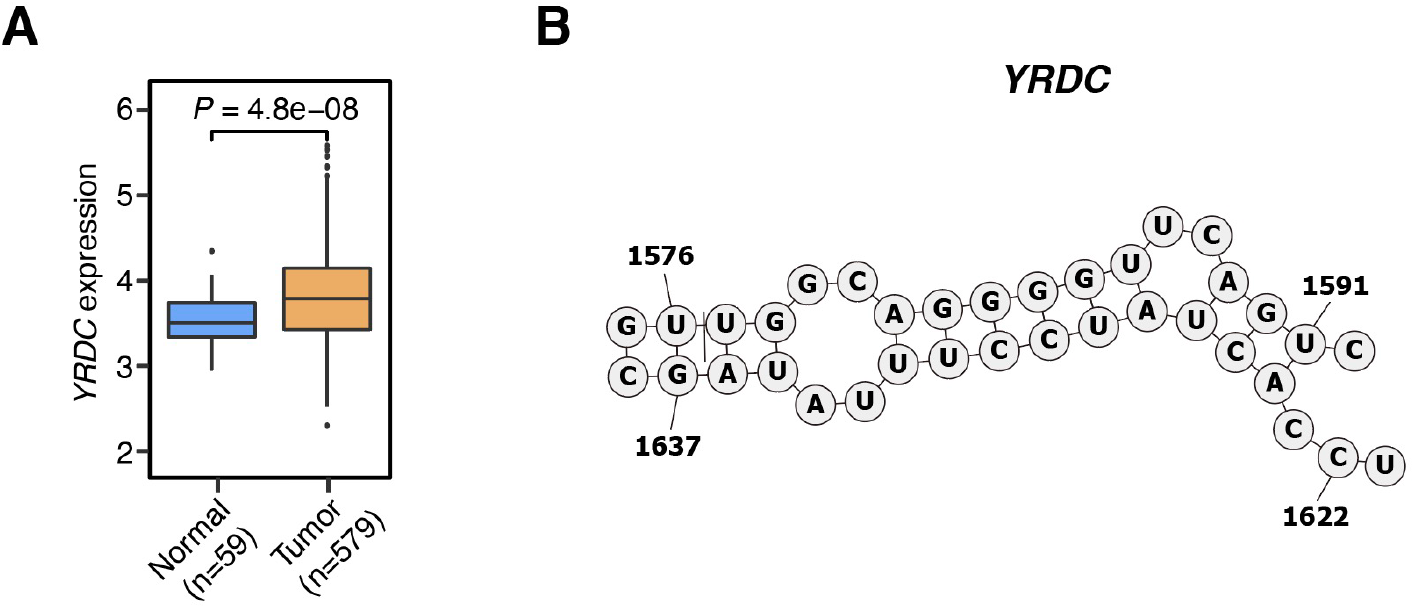
RNA structures is associated with *YRDC* translation regulation in EGFR-TKIs resistance, related to Figure 3. **(A)** Expression of *YRDC* in TCGA lung cancer tissues compared to normal tissues. *P* values were determined by the two-tailed Student’s *t*-test. **(B)** Predicted secondary structure model of the RNA structure in 3’ UTR region in PC9 cells based on PARIS data, annotated with genomic coordinates.

**Figure S5.**
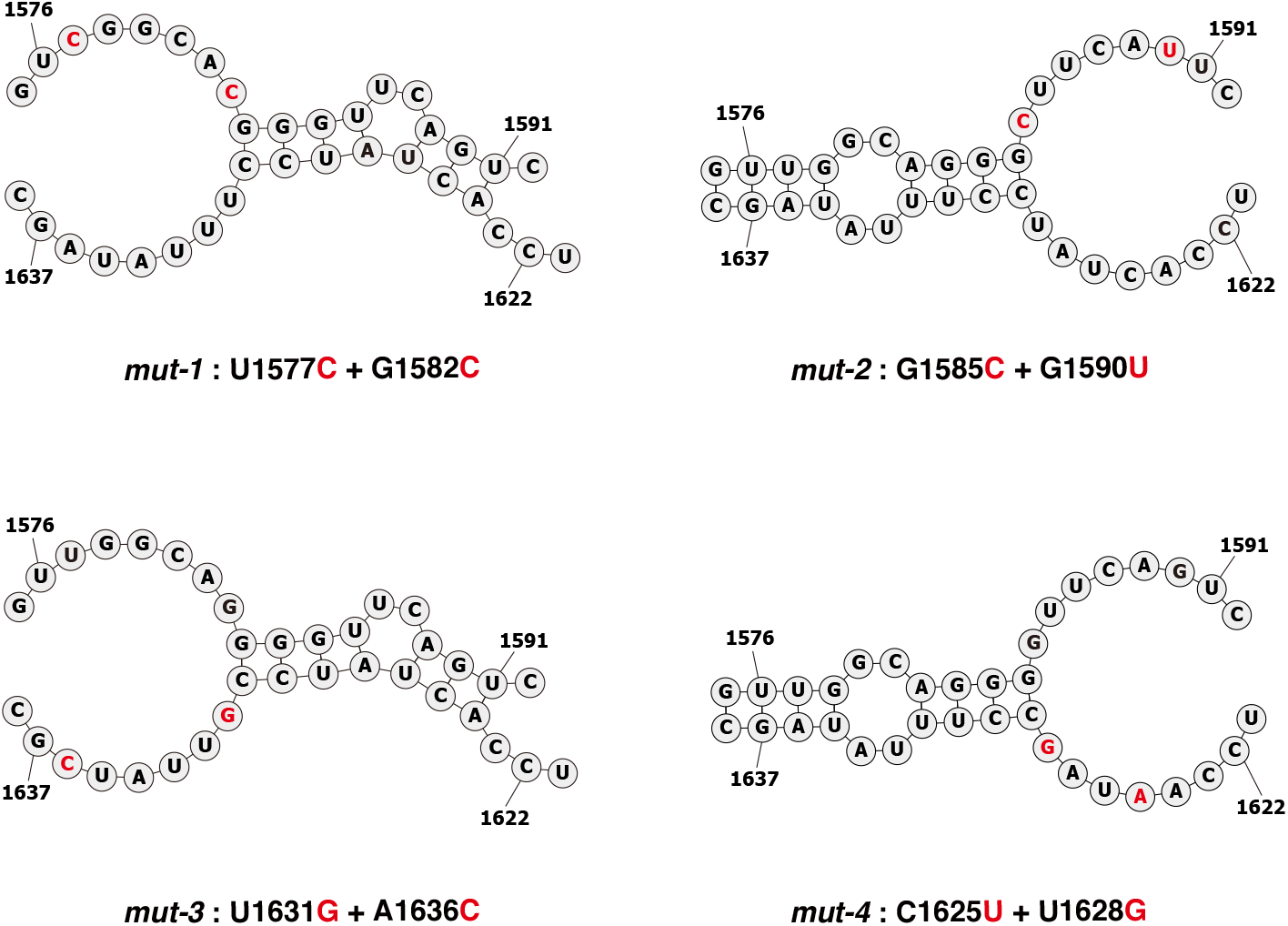
Predicted secondary structure model of the RNA structure in 3’ UTR of designed mutation *YRDC* reporter mRNAs, annotated with genomic coordinates. Red circles represent designs of mutations in the transfection study, related to Figure 4.

**Figure S6.**
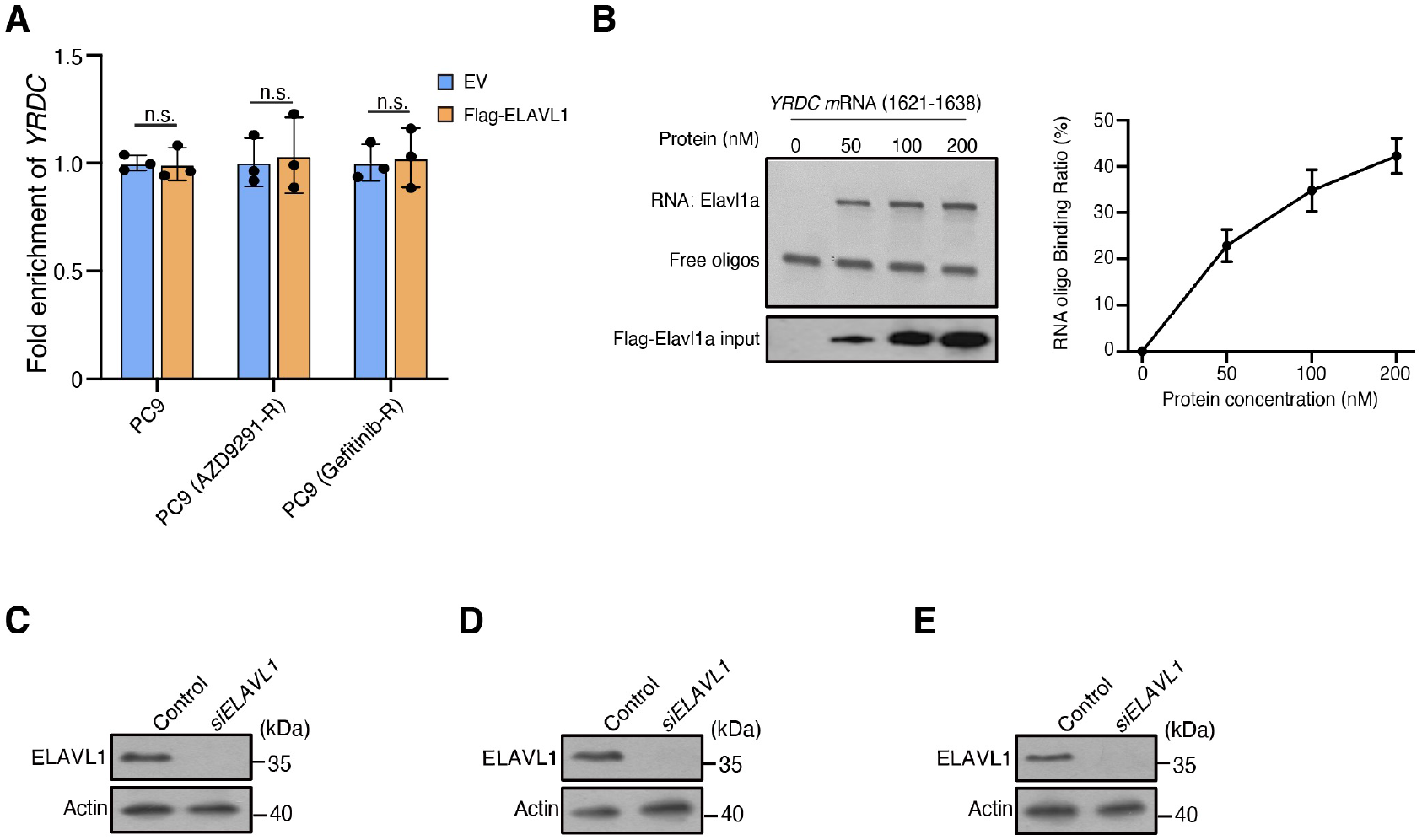
ELAVL1 protein binds 3’UTR of *YRDC* mRNAs, related to Figure 5. **(A)** RIP-qPCR shows the fold enrichment of ELAVL1 binding sites in 3’ UTR of *YRDC* mRNA upon IgG pull-down in PC9 cells, PC9(AZD9291-R) cells and PC9(Gefitinib-R) cells. Error bars, mean ± s.d., n = 3. *P* values were calculated using two-sided Student’s *t*-test, n.s., *P*, not significant. **(B)** EMSA (left) and line graph quantification (right) showing the binding ability of purified Flag-ELAVL1 with RNA probes *(YRDC* 1621-1638). In total, 100 nM of RNA probes was incubated with different concentrations of Flag-ELAVL1 protein. The RNA binding ratio was calculated by (RNA protein) / ((free RNA) + (RNA protein)). Error bars, mean ± s.d., n = 3. **(C)** Western blot demonstrating the absence of ELAVL1 protein upon *siELAVL1* treatment in PC9 cells. **(D)** Western blot demonstrating the absence of ELAVL1 protein upon *siELAVL1* treatment in PC9 (AZD9291-R) cells. **(E)** Western blot demonstrating the absence of ELAVL1 protein upon *siELAVL1* treatment in PC9 (Gefitinib-R) cells.

